# Autism-associated *SCN2A* deficiency disrupts cortico-striatal circuitry in human brain assembloids

**DOI:** 10.1101/2025.06.02.657036

**Authors:** Xiaoling Chen, Jingliang Zhang, Jiaxiang Wu, Morgan J. Robinson, Harish Kothandaraman, Ye-Eun Yoo, Iria M. Gonzalez Dopeso-Reyes, Thomas D. Buffenoir, Manasi S. Halurkar, Zaiyang Zhang, Muhan Wang, Erin N. Creager, Yuanrui Zhao, Maria I. Olivero-Acosta, Kyle W. Wettschurack, Zhefu Que, Chongli Yuan, Allison J. Schaser, Nadia A. Lanman, Jean-Christophe Rochet, William C. Skarnes, Eric J. Kremer, Yang Yang

## Abstract

Profound autism spectrum disorder (ASD) is frequently attributable to single-gene mutations, with *SCN2A* (voltage-gated sodium channel Na_V_1.2) protein-truncating variants (PTVs) being one of the most penetrant. Although cortico-striatal circuitry is implicated as a key node in ASD, the impact of *SCN2A* deficiency on human neural circuits is unknown. Using the human cortico-striatal assembloid model, we show that the autism-causing PTV *SCN2A-C959X* impairs long-range cortical axonal projections, reduces striatal spine density, and attenuates excitatory cortical-striatal synaptic transmission. Surprisingly, these assembloids carrying the heterozygous *SCN2A* nonsense mutation exhibited pronounced network hyperexcitability, a human cell-specific phenotype not observed in *Scn2a^+/-^* mice, highlighting a human-specific circuit vulnerability. Collectively, our study unveils human circuit-specific dysfunctions of *SCN2A* deficiency and *SCN2A*-mediated ASD.

**Highlights:**

- Axonal projections facilitate synapse formation and functional connectivity in human brain assembloids.
- Na_V_1.2 is expressed along neuronal axons, extending to soma and dendrites in human brain assembloids.
- *SCN2A-C959X* disrupts axonal projection patterns, impairs excitatory synaptic transmission, reduces spine density, and results in elevated neuronal excitability.

**Graphical abstract:** 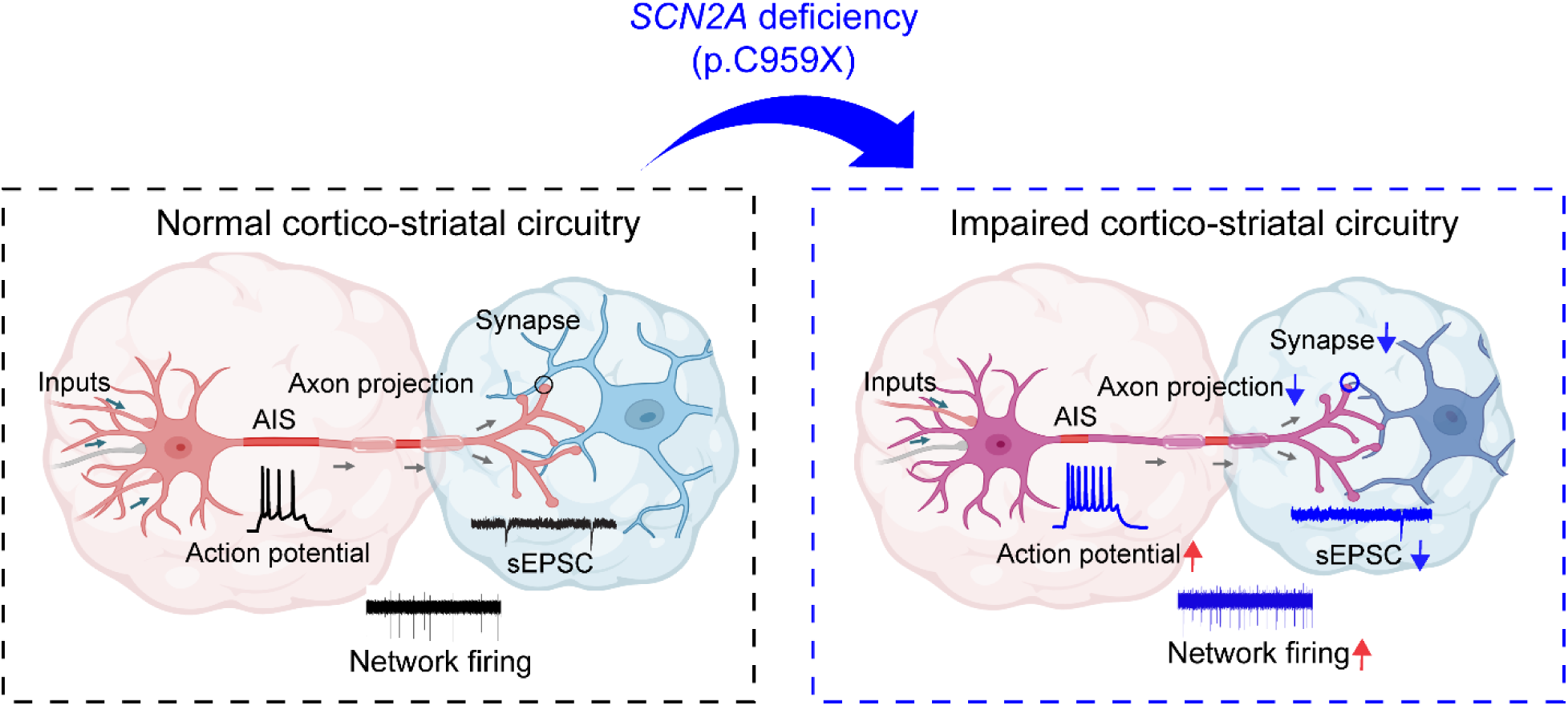

*In brief:* *SCN2A* haploinsufficiency impairs cortico-striatal circuitry.
*SCN2A* haploinsufficiency disrupts axon initial segment (AIS) integrity, leading to hyperexcitability (red arrow), reduced axon projections, and impaired synaptic transmission (decreased sEPSCs and altered network firing). These deficits result in dysfunction within the cortico-striatal circuitry.

## INTRODUCTION

Autism spectrum disorder (ASD) affects about 1 in 31 children in the United States^1^. Although ASD is etiologically heterogeneous^2^, severe forms of the disorder are often caused by mutations in single genes. Among these, loss-of-function (LoF) mutations in *SCN2A*, which encodes the voltage-gated sodium channel Na_V_1.2, have emerged as a leading cause of monogenic ASD^3,4^. In particular, the *de novo* nonsense mutation *SCN2A* c.2877C>A (p.Cys959Ter), referred to as *SCN2A-C959X*, produces a protein-truncating variant (PTV) associated with profound clinical consequences^3,5,6^. Na_V_1.2 is predominantly localized at the axon initial segment (AIS) and plays an essential role in initiating and propagating action potentials in developing neurons. Recent studies in rodent models suggest that Na_V_1.2 also regulates synaptic transmission and dendritic function^7,8^. However, how *SCN2A* deficiency perturbs neuronal communication in the human brain remains largely unknown. Given the growing recognition of ASD as a circuitry disorder, particularly involving cortico-striatal circuitry^9,10^, there is a critical need to elucidate how *SCN2A* deficiency affects this circuit.

The emerging induced pluripotent stem cells (iPSCs)-derived brain organoids have revolutionized the study of human neural development and neurodevelopmental disorders *in vitro*^11,12^. Building on these advances, brain assembloids, which integrate region-specific organoids, offer an unprecedented opportunity to model neuronal connectivity and circuit function across brain regions^13–18^. Here, we leveraged assembloid technology to reconstruct a cortico-striatal network comprising cortical pyramidal neurons and striatal medium spiny neurons. Using this human cell-based model, we systematically explored the consequences of Na_V_1.2 deficiency on human neurons.

Here in this study, we demonstrated that cortical axonal innervation promotes synapse formation and functional connectivity with striatal neurons in assembloids, as revealed by spatial and temporal tracking of axonal projections. By contrast, assembloids carrying the *SCN2A-C959X* mutation exhibited reduced cortical projections, accompanied by decreased dendritic spine density and impaired excitatory synaptic transmission. RNA sequencing further confirmed the downregulation of gene pathways involved in axonal and synaptic development. Unexpectedly, we observed network hyperexcitability of these *SCN2A*-deficient neurons, which is likely attributed to the compensatory adaptation of the impaired synaptic function. Collectively, our findings provide critical insights into the molecular and circuit-level manifestations of *SCN2A* deficiency, demonstrating the utility of human brain assembloids in modeling ASD-associated circuit dysfunction. These results advance our understanding of the role of Na_V_1.2 in neural circuits associated with profound ASD.

## RESULTS

### Cortical projections promote striatal synapse formation and function in brain assembloids

To model the distinct neuronal types and regional identities of the human cortex and striatum *in vitro*, we generated cortical organoids (hCOs) and striatal organoids (hStrOs) from human induced pluripotent stem cells (hiPSCs) following established protocols^13^ (**Figure S1A**). Immunostaining of hCOs confirmed mature neuronal markers neuronal nuclei (NeuN)^+^ and microtubule-associated protein 2 (MAP2)^+^ (**Figure S1B**), as well as cortical layer-specific markers T-box brain transcription factor 1 (TBR1)^+^ and COUP-TF-interacting protein 2 (CTIP2)^+^, with minimal GABAergic neuron marker glutamate decarboxylase1 (GAD67)^+^ (**Figure S1C**), showing the cortical identity. Similarly, hStrOs expressed robust GAD67^+^ and dopamine- and cAMP-regulated phosphoprotein 32 (DARPP32)^+^ (**Figure S1D**), markers for striatal medium spiny neurons (MSNs)^13^, indicative of GABAergic-enriched striatal identity.

To reconstruct the cortico-striatal circuitry, the two types of region-specific organoids were fused to form hCO-hStrO assembloids (**Figure 1A**). Immunostaining confirmed distinct cortical (NeuN^+^/TBR1^+^) and striatal (GABA^+^) compartments (**Figures 1B, C; S1E**). To examine if cortical neurons establish synaptic connectivity with striatal neurons, we implemented anterograde viral tracing by incubating hCOs with AAV1-hSyn1-Cre and, separately, hStrOs with AAV1-Ef1a-DIO-mScarlet and AAV9-hSyn-EGFP. If synaptic connectivity exists, AAV1-hSyn1-Cre would undergo anterograde trafficking into hStrO neurons and express Cre recombinase. Then, Cre activity would flip the DIO-mScarlet cassette and induce mScarlet expression, while EGFP^+^ cells serve as a striatal compartment marker (**Figure 1D**). One month after fusion to form assembloids, we observed cells coexpressing GFP and mScarlet on the hStrOs (**Figure 1D**). These data indicate the presence of cortico-striatal connectivity within the assembloid, which was further confirmed with retrograde labeling (**Figure 1E**). To assess whether the cortico-striatal connections were functional, we conducted chemogenetics coupled with patch-clamp recordings. To evoke presynaptic transmitter release, hCOs were transduced with AAV9-hSyn-Gq-mCherry to express a Gq-DREADD that excites neurons upon CNO administration, while hStrOs were transduced with AAV9-hSyn-EGFP for compartmental distinction. Two months post-fusion, patch-clamp recordings in slices revealed a reversible increase in spontaneous excitatory postsynaptic current (sEPSC) frequency in hStrO neurons following bath application of CNO suggesting functional synaptic connectivity between cortical and striatal neurons (**Figure 1F(i–iv)**).

**Figure 1.**
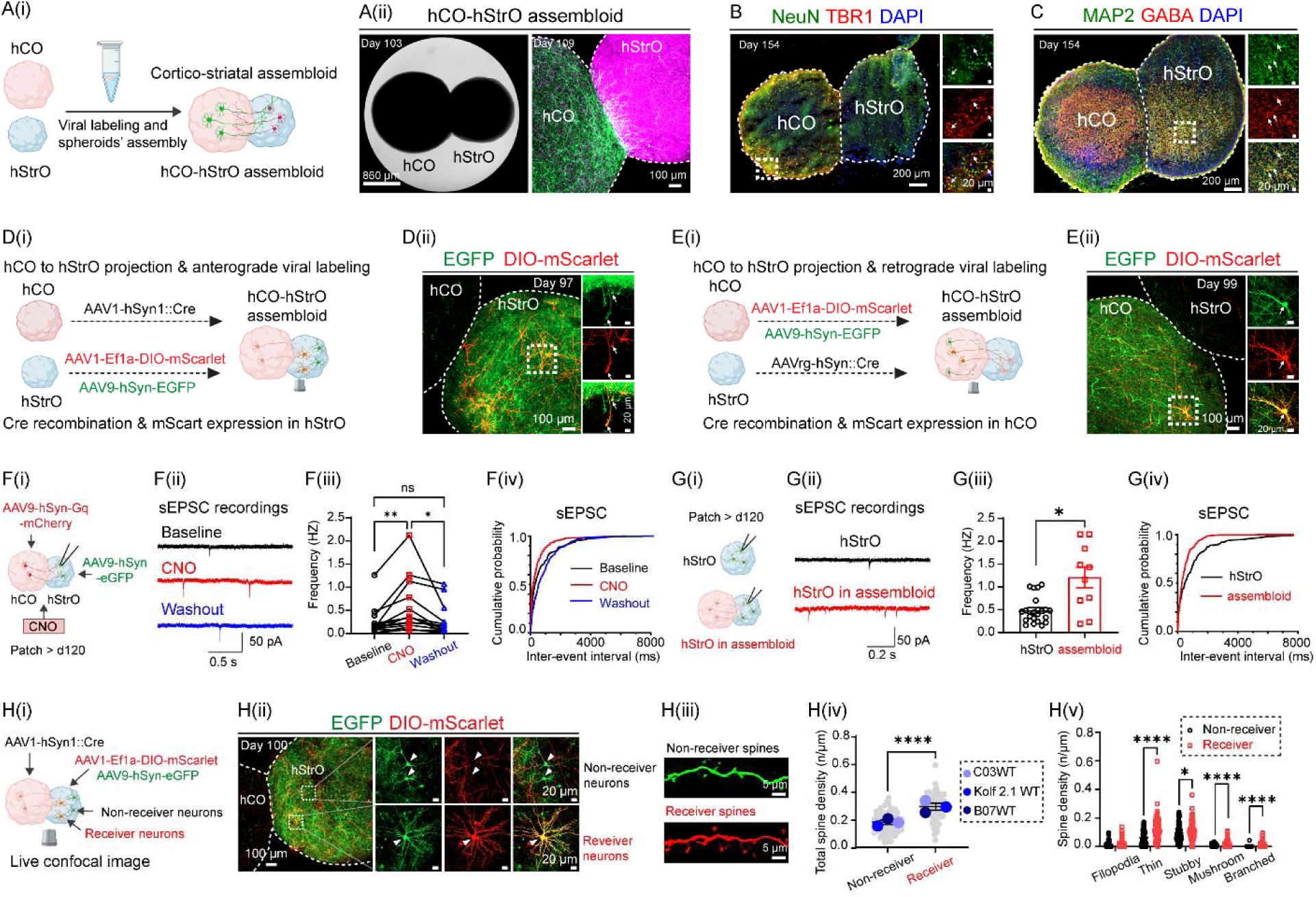
Generation, connectivity mapping, and functional characterization of cortico-striatal assembloids. See also Figure S1. (A) Modeling cortico-striatal assembloids. (A(i)) Schematic depicting the fusion of human cortical (hCO) and striatal organoid (hStrO) to generate cortico-striatal assembloid *in vitro*. (A(ii)) Bright-field image of a fused hCO-hStrO assembloid (left, scale bar: 860 µm) and confocal fluorescence image showing morphological structures of hCO (green) and hStrO (magenta) (right, scale bar: 100 µm). (B, C) Neuronal composition of assembloids. (B) Immunostaining of NeuN (green) and TBR1 (red) on day 154, with a dashed line marking the hCO-hStrO boundary. Insets (right) highlight the colocalization of TBR1^+^/NeuN^+^ in the hCO. DAPI (blue) stains nuclei. Scale bars: 200 µm (main), 20 µm (insets). n = 3 assembloids from two hiPSC lines with similar results. (C) Immunostaining of MAP2 (green) and GABA (red) in an hCO-hStrO assembloid on day 154. The insets (right) highlight GABAergic neurons in the hStrO. n = 3 assembloids from two hiPSC lines with similar results. (D, E) Connectivity mapping using viral tracing. (D) Anterograde tracing. (D(i)) Schematic of anterograde viral tracing strategy. hCOs were transduced with AAV1-hSyn1-Cre, and hStrOs with AAV1-Ef1a-DIO-mScarlet and AAV9-hSyn-EGFP. AAV1-hSyn1-Cre transfers in an anterograde direction and via a transsynaptic mechanism from hCO terminals into hStrO neurons, inducing mScarlet expression, while EGFP serves as a striatal compartment marker. (D(ii)) Confocal images of assembloids showing EGFP^+^ (green) and Cre-dependent mScarlet^+^ (red) in striatal neurons receiving cortical input. n = 3 assembloids with similar results (KOLF2.1J and A11 WT hiPSC lines). (E) Retrograde tracing. (E(i)) Schematic of retrograde labeling using AAVrg-hSyn-Cre. (E(ii)) Confocal image showing hCO neurons labeled with AAV1-Ef1a-DIO-mScarlet and AAV9-hSyn-EGFP (green) projecting into hStrO and receiving retrogradely transported AAVrg-hSyn-Cre, inducing mScarlet expression (red). n = 4 assembloids with similar results (KOLF2.1J and A11 WT hiPSC lines). (F) Chemogenetics activation induces synaptic transmission. (F(i)) Schematic of chemogenetics activation of hCO coupled with patch-clamp recording in hStrO. hCOs were transduced with AAV9-hSyn-Gq-mCherry, while hStrOs were labeled with AAV9-hSyn-EGFP. Whole-cell patch-clamp recordings were performed on hStrO neurons, with CNO applied to activate Gq signaling in hCO. (F(ii)) Representative spontaneous excitatory postsynaptic currents (sEPSC) traces in hStrO neurons at baseline (black), CNO (red), and washout (blue) conditions. (F(iii)) sEPSC frequency quantification and (Fiv) cumulative probability of sEPSC inter-event intervals. (n = 16 cells from 6 assembloids, 3 WT cell lines: C03, A11, KOLF2.1J). Mixed-effects analysis with Geisser-Greenhouse correction, Tukey’s multiple comparisons test. ns: not significant, *p < 0.05, **p < 0.01. (G) sEPSC recordings comparing hStrO alone vs. hCO-hStrO assembloids. (G(i)) Schematic of sEPSC recordings from hStrO alone (black) or hStrO in assembloids (red). (G(ii)) Representative sEPSC traces and (G(iii)) quantification of sEPSC frequency (n = 22 cells from 3 hStrO, n = 10 cells from 3 hCO-hStrO assembloids). Unpaired Welch’s t-test: p < 0.05. (G(iv)) Cumulative probability plot of sEPSC inter-event intervals. (H) Live confocal imaging of dendritic spine analysis. (H(i)) Schematic of receiver vs. non-receiver neurons in hStrO. hCOs were transduced with AAV1-hSyn1-Cre, while hStrOs were labeled with AAV1-Ef1a-DIO-mScarlet and AAV9-hSyn-EGFP. (H(ii)) Confocal images showing non-receiver (EGFP^+^ only) and receiver (EGFP^+^/mScarlet^+^, yellow) hStrO neurons. (H(iii)) Representative dendritic spines from non-receiver (green, top) and receiver (red, bottom) hStrO neurons. (H(iv)) Total spine density quantification, comparing non-receiver (gray, n = 69 dendrites) vs. receiver neurons (red, n = 81 dendrites) from 9 hCO-hStrO assembloids across three WT hiPSC lines (C03, KOLF2.1J, B07). ****p < 0.0001 (Mann-Whitney test). (H(v)) Spine subtype classification (filopodia, thin, stubby, mushroom, branched). *p < 0.001, ****p < 0.0001 (Multiple Mann-Whitney test). Data are represented as mean ± SEM.

Cortical glutamatergic projections are known to be instrumental for striatal MSN maturation^19^. To examine how cortical projections influence striatal synaptic properties, we compared sEPSC frequency in striatal neurons within cortico-striatal assembloids versus non-fused hStrO (**Figure 1G(i)**). Two months post-fusion, we observed a higher frequency of sEPSCs within hCO-hStrO than in hStrOs alone at the same *in vitro* development stage (**Figure 1G(ii–iv)**), suggesting that connections with hCOs enhance striatal synaptogenesis. Moreover, to determine whether increased synapse formation correlated with cortico-striatal connectivity, we distinguished striatal neurons receiving direct cortical input (“receiver”) from those without cortical input (“non-receiver”) in assembloids with dual viral labeling and examined their excitatory input. Specifically, hCOs were labeled with AAV1-hSyn1-Cre, while hStrOs were labeled with AAV1-Ef1a-DIO-mScarlet and AAV9-hSyn-EGFP (**Figure 1H(i)**). Receiver neurons in hStrO (EGFP^+^/mScarlet^+^) exhibited significantly higher dendritic spine density and a greater proportion of more mature spine types than non-receiver neurons (EGFP^+^ only) (**Figure 1H(ii– v)**). Together, these results demonstrate that cortico-striatal assembloids derived from hiPSCs successfully establish functional connectivity that promotes neuronal maturation, providing a robust *in vitro* model for studying circuit-level interactions in neurodevelopmental disorders.

### *SCN2A-C959X* mutation disrupts cortico-striatal circuit in brain assembloids

*Scn2a* is robustly expressed in the cortico-striatal circuitry^20^, which is critically involved in ASD^9^. To investigate how ASD-associated *SCN2A* deficiency affects neuronal function in this circuitry, we used CRISPR/Cas9 genome editing to introduce the *SCN2A* c.2877C>A (p.Cys959Ter) mutation (identified in children with profound autism) into reference KOLF2.1J iPSCs (**Figures 2A; S2A**). Sanger sequencing confirmed the successful generation of heterozygous (HET) and homozygous (HOM) mutant lines as well as isogenic wild-type (WT) controls (**Figure 2A**). Western blot analysis revealed significantly reduced Na_V_1.2 protein levels in HET and HOM hCOs compared with WT (**Figure 2B**). qPCR analysis further showed a dose-dependent decrease in *SCN2A* mRNA levels in hCO-hStrO assembloids (**Figure 2C**), consistent with a LoF/deficiency phenotype of this mutation. Immunohistology assays using murine brains localize the Na_V_1.2 signal on the axon initial segment (AIS), where action potential (AP) initiates^7^. Our immunostaining demonstrated that Na_V_1.2 colocalized with Ankyrin-G, a marker of the AIS (**Figure 2D**), indicating that the localization of the Na_V_1.2 channel in both hCO and hStrO neurons is consistent with the murine brain studies. To determine how *SCN2A* deficiency impacts the AIS, we quantified the AIS length across genotypes. Interestingly, we found that HET *SCN2A-C959X* neurons displayed shorter AIS lengths in hCO slices (**Figure S2B(i, ii)**), suggesting that *SCN2A* deficiency damages AIS plasticity, which is critically involved in neuronal intrinsic excitability^21^. Surprisingly, whole-cell patch-clamp recordings revealed that *SCN2A*-deficient neurons exhibited enhanced excitability relative to WT controls, characterized by a shift in the input-output relationship of injected current to AP firing (**Figure 2E(i, ii)**). This neuronal hyperexcitability was further evidenced by elevated maximum firing frequency, increased input resistance, and reduced current rheobase for AP spiking (**Figure 2E(iii–v)**).

**Figure 2.**
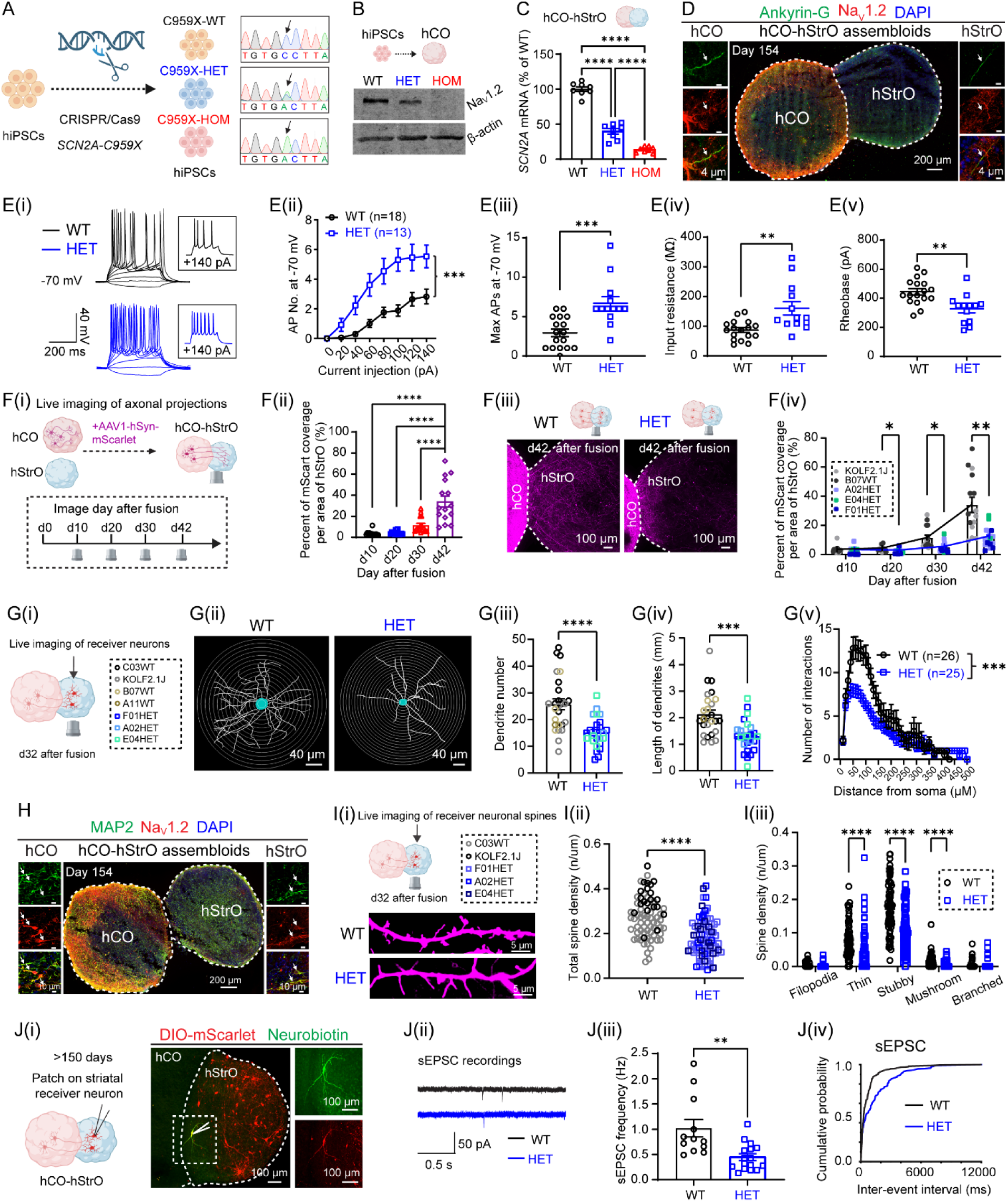
Human assembloids carrying the *SCN2A-C959X* mutation exhibit impaired cortico-striatal circuits. See also Figure S2. (A) CRISPR/Cas9 editing introduced the *SCN2A* p.C959X mutation in human iPSCs. Sanger sequencing confirms successful editing: WT (top) retains cytosine (C), HET (middle) shows overlapping peaks and HOM (bottom) displays a complete C-to-A substitution, introducing a premature stop codon (X). (B) Western blot showing Na_V_1.2 (*SCN2A*) expression in WT, HET, and HOM hCOs, with β-actin as a reference. (C) *SCN2A* mRNA levels in hCO-hStrO assembloids showing reduced expression in HET and HOM *SCN2A-C959X* mutants vs. WT. n = 8 assembloids/group from 3 hiPSC lines (WT: KOLF2.1J, B07, C03; HET: A02, E04, F01; HOM: A03, D06, F03). One-way ANOVA with Tukey’s test, ****p < 0.0001. (D) Immunostaining of Ankyrin-G (green), Na_V_1.2 (red) in hCO-hStrO assembloids on day 154, demonstrating Na_V_1.2 localization at the axon initial segment (AIS). n = 3 assembloids. (E) Electrophysiological alterations in neurons carrying *SCN2A-C959X* mutation. (E(i)) The representative traces of action potentials (APs) in WT and HET at −70 mV. Inset: AP response to +140 pA current injection. (E(ii)) AP generated in response to depolarizing current pulses at −70 mV. (**p < 0.01, two-way ANOVA). (E(iii)) Individual and average maximum AP counts at −70 mV. Mann-Whitney test, ***p < 0.0001. (E(iv)) Input resistance and (E(v)) rheobase measurements at −70 mV. Unpaired t-test. **p < 0.001. (F) Axon projection deficits in *SCN2A-C959X* assembloids. (F(i)) Experimental timeline and (F(ii)) quantification of axon projections from hCO to hStrO in assembloids on days 10, 20, 30, and 42 post-fusion. One-way ANOVA, Tukey’s test, ****p < 0.0001. (F(iii)) Representative images of axon projections (mScarlet⁺) on day 42 in WT and HET assembloids. (F(iv)) mScarlet coverage quantification in hStrO on days 10, 20, 30, and 42, comparing WT (n = 15 assembloids from B07, KOLF2.1J) and HET (n = 16 assembloids from F01, A02, E04). Mixed-effects model with Geisser-Greenhouse correction, Sidak’s test. *p < 0.05, **p < 0.01. (G) Dendritic morphology deficits in *SCN2A-C959X* assembloids. (G(i)) Schematic of receiver neuron identification in hStrO. (G(ii)) Representative images of neuronal morphology (Sholl analysis) in WT and HET. (G(iii)) Total dendrite number (unpaired t-test), (G(iv)) dendrite length (unpaired t-test), and (Gv) interaction number (two-way ANOVA) in WT (n = 26 neurons, B07, C03, A11, KOLF2.1J cell lines) and HET (n = 25 neurons, F01, A02, E04 cell lines). ***p < 0.001, ****p < 0.0001. (H) Na_V_1.2 localization in dendrites. Immunostaining for MAP2 (green), Na_V_1.2 (red), and DAPI (blue) in hCO-hStrO assembloids, confirming Na_V_1.2 expression in neuronal dendrites. n = 3 assembloids from two hiPSC lines with consistent results. (I) Dendritic spine deficits in *SCN2A-C959X* assembloids. (I(i)) Confocal images of striatal receiver (mScarlet⁺) dendritic spines in WT and HET. (I(ii)) Quantification of total receiver spine density in WT (gray, n = 74 dendrites, 9 assembloids, C03, KOLF2.1J, B07 lines) and HET (blue, n = 86 dendrites, 9 assembloids, F01, A02, E04 lines). Mann-Whitney test, ****p < 0.0001. (I(iii)) Spine subtype analysis. ****p < 0.0001 (Multiple Mann-Whitney test). (J) Synaptic transmission deficits in *SCN2A-C959X* assembloids. (J(i)) Confocal images of receiver neurons (mScarlet⁺, red) in hStrO from hCO-hStrO assembloids. Patch-clamped neurons labeled with neurobiotin (green) confirm that recorded neurons are receivers. (J(ii)) Example sEPSC traces in hStrO receiver neurons from WT (black, top) and HET (blue, bottom). (J(iii)) sEPSC frequency in WT (n = 12 cells, 7 assembloids, A11, KOLF2.1J lines) and HET (n = 16 cells, 7 assembloids, A02, E04 lines). Unpaired t-test, **p < 0.01. (J(iv)) Cumulative probability of sEPSC inter-event intervals. Data are represented as mean ± SEM.

To assess the consequences of *SCN2A* deficiency on brain circuitry beyond neuronal intrinsic properties, we examined connectivity in cortico-striatal assembloids. Using assembloids prepared from WT organoids, we observed a progressive increase in mScarlet^+^ projections from hCOs into hStrOs after fusion (**Figure 2F(i, ii)**). In contrast, HET *SCN2A-C959X* assembloids exhibited significantly reduced axonal innervation into hStrOs compared with WT on days 20, 30, and 42 after fusion (**Figure 2F(iii, iv)**), indicating that *SCN2A* deficiency impairs long-range inter-brain regional projections. Given the potential impact of impaired cortical projections on neuronal maturation, we further analyzed the morphology of hStrO receiver neurons innervated by cortical input using dual AAV labeling. hCOs and hStrOs were transduced with AAV1-hSyn1-Cre or AAV1-Ef1a-DIO-mScarlet, respectively. One month after fusion, Sholl analysis of HET receiver neurons (EGFP^+^/mScarlet^+^) in the hStrO compartment revealed significantly reduced dendritic complexity compared with their WT counterparts, including fewer dendritic branches, shorter dendritic length, and decreased interaction points (**Figure 2G(i–v)**). These findings link decreased Na_V_1.2 expression to impaired axonal innervation and neuronal maturation within assembloids.

Previous results from mice also suggest that Na_V_1.2 localizes to soma and dendrites^8^. Consistently, we found Na_V_1.2 is colocalized with neuronal cell bodies (NeuN^+^) and neurites (MAP2^+^) in both cortical and striatal compartments of human assembloids (**Figures 2H, S2D**), suggesting a role of Na_V_1.2 in soma and dendrites. To assess the influence of *SCN2A-C959X* on excitatory synapses, we first confirmed that cortical axons can form excitatory synapses by labeling presynaptic terminals with Syn1 and postsynaptic sites with PSD95. In contrast to controls, quantitative analysis of Syn1^+^/PSD95^+^ immunoreactivity revealed a significant reduction in their colocalization in HET *SCN2A-C959X* neurons, indicating impaired excitatory synaptic connectivity (**Figure S2C(i–iii)**). Because dendritic spines serve as the primary sites for excitatory synapses, we next characterized spine morphology in live hCO-hStrO assembloids. HET receiver neurons (mScarlet^+^) exhibited a markedly reduced spine density compared with WT, with thin, stubby, and mature mushroom spines being particularly affected (**Figure 2I(i–iii)**), indicating structural deficits of dendritic spines. To determine whether these morphological changes translate into functional impairments, we performed patch-clamp recordings on receiver neurons (mScarlet^+^) in hCO-hStrO assembloid slices (**Figure 2J(i)**). HET receiver neurons displayed a significant decrease in sEPSC frequency relative to WT (**Figure 2J(ii–iv**)), indicating disrupted synaptic transmission. *Post hoc* immunostaining of recorded neurons further confirmed lower dendritic spine density (**Figure S2E(i–v)**), consistent with our live-cell imaging results (**Figure 2I(i–iii)**).

To further enhance the rigor of our findings, we also examined assembloids carrying the HOM *SCN2A-C959X* mutation and observed similar phenotypes, including reduced dendritic complexity and reduced spine density (data not shown). Collectively, our data demonstrate that *SCN2A-C959X* mutation disrupts AIS plasticity, axonal projections, dendritic architecture, and synaptic function in human cortico-striatal assembloids. These comprehensive phenotypes underscore the critical role of Na_V_1.2 in establishing and maintaining functional connectivity within cortico-striatal circuits, providing insights into how *SCN2A* LoF variants contribute to the pathology of ASD.

### *SCN2A-C959X* mutation impairs axon- and synapse-related gene pathways

To elucidate the molecular basis underlying *SCN2A* deficiency in the cortico-striatal circuitry, we conducted bulk RNA sequencing on hCO-hStrO assembloids across genotypes (**Figure 3A**). Principal component analysis (PCA) demonstrated distinct clustering among WT, HET, and HOM groups, highlighting global shifts in gene expression profiles (**Figure S3A**). We verified the *SCN2A* expression levels, which were largely reduced in both HET and HOM groups (**Figure 3B**), consistent with the qPCR results shown in **Figure 2C**. We then identified 6,145 differentially expressed genes (DEGs) in HET vs. WT (2,846 upregulated; 3,299 downregulated; **Figure 3C(i)**), and 8,907 DEGs in HOM vs. WT (4,717 upregulated; 4,190 downregulated; **Figure 3C(ii)**). Our subsequent analysis prioritized overlapping DEGs shared between HET and HOM groups, as these likely represent core pathogenic mechanisms of *SCN2A* deficiency. Since we found that *SCN2A-C959X* affects neuronal excitability and synaptic transmission (**Figure 2E, J**), we explored the genes associated with these functions. Indeed, our analysis revealed significant downregulation of critical genes related to voltage-gated sodium channels (*SCN2A, SCN8A, SCN1A*), axonal integrity (*ANK2, ANK3*), and synaptic pathways (*SYN1, DLG4, GRIA2, GRIA3, GRIA4, GRIN1, GRIN2A, GRIN3A*) (**Figure 3C(i, ii)**).

**Figure 3.**
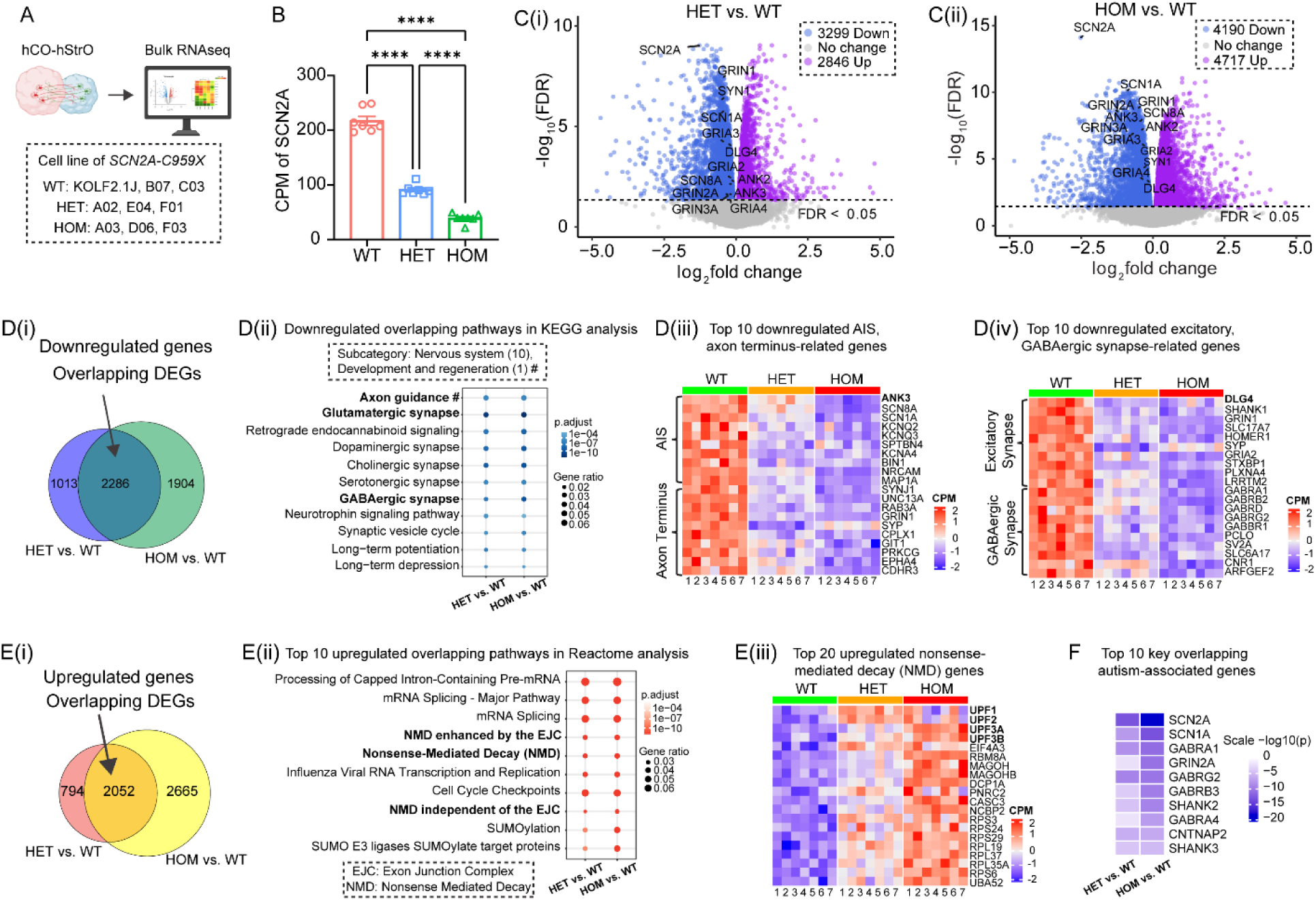
Transcriptomic analysis reveals differential gene expression and pathway alterations in assembloids carrying *SCN2A-C959X* mutation. See also Figure S3. (A) Experimental workflow. n = 7 assembloids/group from WT (iPSC lines: KOLF2.1J, B07, C03), HET (iPSC lines: A02, E04, F01), and HOM (iPSC lines: A03, D06, F03). (B) Bar graph showing counts per million (CPM) of *SCN2A* expression in WT, HET, and HOM in bulk RNA sequencing. One-way ANOVA with Tukey’s comparisons, ****p < 0.0001. (C) Differential gene expression analysis. Volcano plots displaying differentially expressed genes (DEGs) in HET vs. WT (C(i)) and HOM vs. WT (C(ii)) comparisons. Upregulated genes (magenta), downregulated (blue), and non-significant (gray) (FDR < 0.05). Key downregulated genes associated with axon and synapse pathways are labeled. (D) Downregulated gene pathways. (D(i)) Venn diagram illustrating overlapping downregulated genes in HET vs. WT and HOM vs. WT. (D(ii)) KEGG pathway enrichment analysis of downregulated genes in the nervous system and development and regeneration (marked with ’#’) classification. (D(iii)) Axonal deficits. Heatmap of top downregulated axon-related genes, categorized into axon initial segments (AIS) (top 10) and axon terminal genes (top 10), indicating axonal deficits in HET and HOM. (D(iv)) Synaptic deficits. Heatmap of top downregulated synapse-related genes, classified into excitatory (top 10) and inhibitory (GABAergic, top 10) synapses. (E) Upregulated gene pathways. (E(i)) Venn diagram showing overlapping upregulated genes in HET vs. WT and HOM vs. WT. (E(ii)) Reactome pathway enrichment analysis of the top 10 overlapping upregulated pathways, with key pathways associated with nonsense-mediated decay (NMD) highlighted in bold. (E(iii)) Heatmap of top 20 upregulated NMD-related genes in HET and HOM vs. WT (CPM: Counts per million). (F) Autism-associated gene alterations. Heatmap of the top 10 ASD-associated DEGs in HET vs. WT and HOM vs. WT. Scale: -log_10_(P).

Further investigation of overlapping DEGs identified 2,286 consistently downregulated genes (**Figure 3D(i)**). KEGG pathway enrichment analysis of these genes revealed significant disruptions in pathways related to neural development, axon guidance, and synaptic signaling (**Figure 3D(ii)**). GO cellular component (CC) enrichment further reinforced deficits in axon terminals, AIS, excitatory synapses, and GABAergic synapses (**Figures 3D(iii, iv); S3B**). The observed downregulation of *ANK3* (ankyrin-G) and DLG4 (PSD-95) is in alignment with protein-level changes in our immunostaining experiments (**Figure S2B, C**). Additionally, GO molecular function (MF) and Reactome analyses confirmed the primary impact of *SCN2A* deficiency on synaptic pathways (**Figure S3C, E**). Moreover, GO biological process (BP) analysis highlighted pronounced downregulation in multiple ion channel categories, specifically, ligand-gated channels, and voltage-gated sodium and potassium channels (**Figure S3D(i–iv)**). This dysregulation potentially could explain the altered neuronal excitability seen in *SCN2A-C959X* mutant neurons.

Conversely, examination of overlapping upregulated genes (2,052 genes; **Figure 3E(i)**) using Reactome pathway analysis highlighted enrichment in processes including nonsense-mediated decay (NMD), mRNA processing, mRNA splicing, and exon junction complex (EJC) activities (**Figures 3E(ii, iii); S3F**), indicative of enhanced RNA surveillance mechanisms. Reactome was selected for its comprehensive annotations of RNA metabolism pathways, which are less extensively represented in other databases such as KEGG. Given our western blot and qPCR results demonstrating substantial reductions in *SCN2A* mRNA and protein levels (**Figure 2B, C**), these molecular profiling findings indicate that the C959X mutation likely drives decreased *SCN2A* expression via activation of NMD-related pathways.

Interestingly, network analysis revealed 59 autism-related DEGs that were consistently altered in both HET and HOM assembloids (**Figure S3G**). The heatmap of the top 10 autism-related genes (**Figure 3F**) illustrates their correlative expression patterns, providing additional insights into shared neurobiological mechanisms by which distinct genetic risk factors may converge to drive ASD pathology^22^. Collectively, our transcriptome analysis demonstrates that *SCN2A-C959X* prominently disrupts axonal and synaptic pathways, impairing synaptic connectivity and neuronal function in human cortico-striatal circuits. Such dysregulation may underlie ASD-associated neural dysfunction, providing molecular insights into the impact of *SCN2A* LoF variants on ASD etiology.

## DISCUSSION

ASD is a complex neurodevelopmental condition, and identifying molecular perturbations caused by monogenic mutations such as those in the *SCN2A* gene provides valuable insights into pathophysiology. Here, we used human cortico-striatal assembloids to investigate how the *SCN2A-C959X* mutation disrupts neuronal function and circuitry connectivity. Our findings demonstrate that *SCN2A* deficiency leads to pronounced impairments in axonal projections, dendritic spine morphology, synaptic transmission, and neuronal excitability, underscoring the essential role of Na_V_1.2 in establishing and maintaining functional cortico-striatal circuitry.

Na_V_1.2 plays multiple roles in neurons: it extends from the AIS to middle and distal axons, determining action potential (AP) initiation and propagation, while also localizing to soma and dendrites regulating AP backpropagation^7,8^. Consistently, our immunostaining localizes Na_V_1.2 with Ankyrin-G at the AIS, NeuN in the soma, and MAP2 in dendrites in human assembloids (**Figure 2**). In the pathological condition, *SCN2A* deficiency in axons likely attenuates AP peaks during propagation, thus may lead to reduced neurotransmitter release and impaired axonal projections and synaptic connectivity. We observed that *SCN2A-C959X* disrupts cortical axonal projections into the striatum, leading to reduced dendritic spine density, altered morphology, and impaired synaptic transmission in striatal receiver MSNs (**Figure 2**). These findings align with rodent studies demonstrating that Na_V_1.2 dysfunction impairs AP backpropagation, ultimately leading to defective synaptic function^7,8^. Additionally, we found that *SCN2A-C959X* results in AIS shortening, a notable finding given that longer AIS length is typically associated with increased neuronal intrinsic excitability, as increased Na_V_ conductance in the longer AIS lowers the AP voltage threshold^21^. Paradoxically, despite a shortened AIS, *SCN2A*-deficient cortical neurons in our model exhibited intrinsic hyperexcitability (**Figure 2**). Interestingly, this counterintuitive hyperexcitability coincided with downregulated potassium channels (**Figure S3**), which is surprisingly consistent with our previous observations from *Scn2a*-deficient mice^20^. We proposed that potassium channel downregulation could be a compensatory mechanism to counterbalance Na_V_1.2 loss, ultimately leading to neuronal hyperexcitability^20^.

It is noteworthy that the neuronal hyperexcitability caused by a heterozygous *SCN2A* nonsense mutation (∼50% reduction of *SCN2A* expression) is a human-cell-specific phenotype. Mouse models require over 70% reduction in *Scn2a* expression to exhibit similar hyperexcitability^20,23^, underlining critical differences between human and mouse cellular physiology and emphasizing the necessity of utilizing human cell models to study human disorders. Another advantage of using organoid and assembloid models is their ability to mature over extended periods (hundreds of days), enabling neurons to develop complex architectures^24^ and electrophysiological properties^25^ closely resembling physiological conditions^26^. These advanced features may explain the observed discrepancy between our findings and previous studies using *SCN2A^+/-^* hiPSC-derived neurons cultured in 2D. Specifically, 2D cultured *SCN2A^+/-^* neurons exhibit increased AIS length and reduced neuronal excitability^27^, unlike what we reported in this study (**Figure 2**). It is known that neurons cultured in 2D monolayers represent relatively immature stages^28^. In contrast, the enhanced maturation of 3D assembloids may uncover phenotypes not detectable at earlier developmental stages.

Principal neurons in the cortico-striatal circuitry typically do not fire spontaneously (**Figures 2E(i, ii); 4E(i, ii)**), yet *SCN2A-C959X* neurons are intrinsically hyperexcitable (**Figure 2**). Despite impaired excitatory transmission, MEA recordings revealed an increased neuronal firing rate in *SCN2A-C959X* assembloids (**Figure 4**). This striking reversible network-level hyperexcitability may arise from the compensatory synaptic changes (e.g., possible imbalance of excitatory and inhibitory synapses; **Figures 3; S3**), consistent with our previous finding in *Scn2a*-deficient mice, which show both synaptic impairments and enhanced neuronal firing *in vivo*^29^. This raises intriguing questions: **1)** are circuit-level deficits, including axonal projection, spine density, and synaptic neurotransmission, more closely linked to ASD pathogenesis than intrinsic excitability alone? **2)** does prolonged impairment in neuronal communication trigger compensatory intrinsic hyperexcitability to counterbalance reduced neurotransmitter release? Addressing these questions in future studies will further elucidate the complexity of neuronal excitability regulation in disease pathology.

*SCN2A* mutations in patients are heterozygous. The *SCN2A-C959X* nonsense mutation leads to a premature stop codon, likely triggering nonsense-mediated decay^6,30^ and resulting in haploinsufficiency. In this condition, the 50% expression (**Figure 2**) from the unaffected allele is insufficient to maintain normal neuronal function. Our RNA-seq data revealed an upregulation of NMD pathways (**Figure 3**), further supporting haploinsufficiency as the primary pathogenic mechanism. By contrast, *SCN2A* LoF mutations particularly missense mutations could also result in a dominant-negative effect^31–33^, producing mutant proteins that aggregate and sequester WT Na_V_1.2, further reducing functional Na_V_1.2 levels. While our homozygous *SCN2A-C959X* model does not mimic the heterozygous state found in patients, it represents an extreme form of *SCN2A* deficiency, likely revealing important biological insights related to *SCN2A* functions in human neurons. Together, our results demonstrate that human brain assembloids carrying both HET and HOM *SCN2A* mutation could serve as a powerful platform for studying the cellular and circuitry pathology of ASD.

In summary, our study demonstrates that the autism-causing *SCN2A-C959X* profoundly disrupts cortico-striatal connectivity in human brain assembloids, leading to synaptic deficits, altered gene expression, and network hyperexcitability. Conceptually, our work significantly advances beyond prior *Scn2a* mouse model studies by directly linking species-specific neural circuitry disruptions with functional outcomes relevant to human pathology. More broadly, these findings underscore the power of human brain organoids/assembloids in modeling neurodevelopmental disorders and the potential of advancing precision medicine approaches, laying a foundational framework applicable to other monogenic ASDs.

## RESOURCE AVAILABILITY

### Lead contact

Further information and requests for resources should be directed to and will be fulfilled by the lead contact, Yang Yang.

### Materials availability

This study did not generate new unique reagents.

### Data and code availability

Detailed datasets and source code supporting the current study are available from the corresponding author on request.

## ACKNOWLEDGMENTS

Research reported in this publication was partially supported by the National Institute of Neurological Disorders and Stroke of the National Institutes of Health under Award Number R01NS117585 and R01NS123154 to Y.Y. The authors gratefully acknowledge support from the Purdue Institute for Drug Discovery and the Purdue Institute for Integrative Neuroscience for additional funding support. X.C. was supported by the AES Postdoctoral Research Fellowship. The Yang lab is grateful to the *FamilieSCN2A* Foundation for the Hodgkin-Huxley Research Award to Y.Y. and the Action Potential Grant support to X.C., J.Z., and Y.E.Y. The Yang lab appreciates the bioinformatics support of the Collaborative Core for Cancer Bioinformatics (C^3^B) with support from the Indiana University Simon Comprehensive Cancer Center (Grant P30CA082709), Purdue Institute for Cancer Research (Grant P30CA023168), and Walther Cancer Foundation. The content is solely the responsibility of the authors and does not necessarily represent the official views of the Indiana State Department of Health or the National Institutes of Health.

## AUTHOR CONTRIBUTIONS

X.C., J.Z., and Y.Y. designed the experiments. X.C., J.Z., M.J.R., M.S.H., Z.Z., M.W., and E.N.C. performed the experiments. W.C.S. designed and performed the *SCN2A* gene editing experiment. I.G.D.R., T.B., and E.J.K. generated and provided unpublished reagents. J.W., H.K., Y.-E.Y., Y.Z., M.I.O.A., K.W.W., Z.Q., C.Y., A.J.S., N.A.L., J.-C.R., and W.C.S. participated in data analysis and experimental design. Y.Y. supervised the project. X.C., J.Z., and Y.Y. wrote the paper with input from all authors.

## DECLARATION OF GENERATIVE AI AND AI-ASSISTED TECHNOLOGIES IN THE WRITING PROCESS

During the preparation of this work, the authors used ChatGPT 4.5 to improve the readability and language in this manuscript while ensuring that the main conclusions remained unchanged. After using this tool, the authors reviewed and edited the wording as necessary and take full responsibility for the content of the publication.

## SUPPLEMENTAL INFORMATION

**Figure S1.**
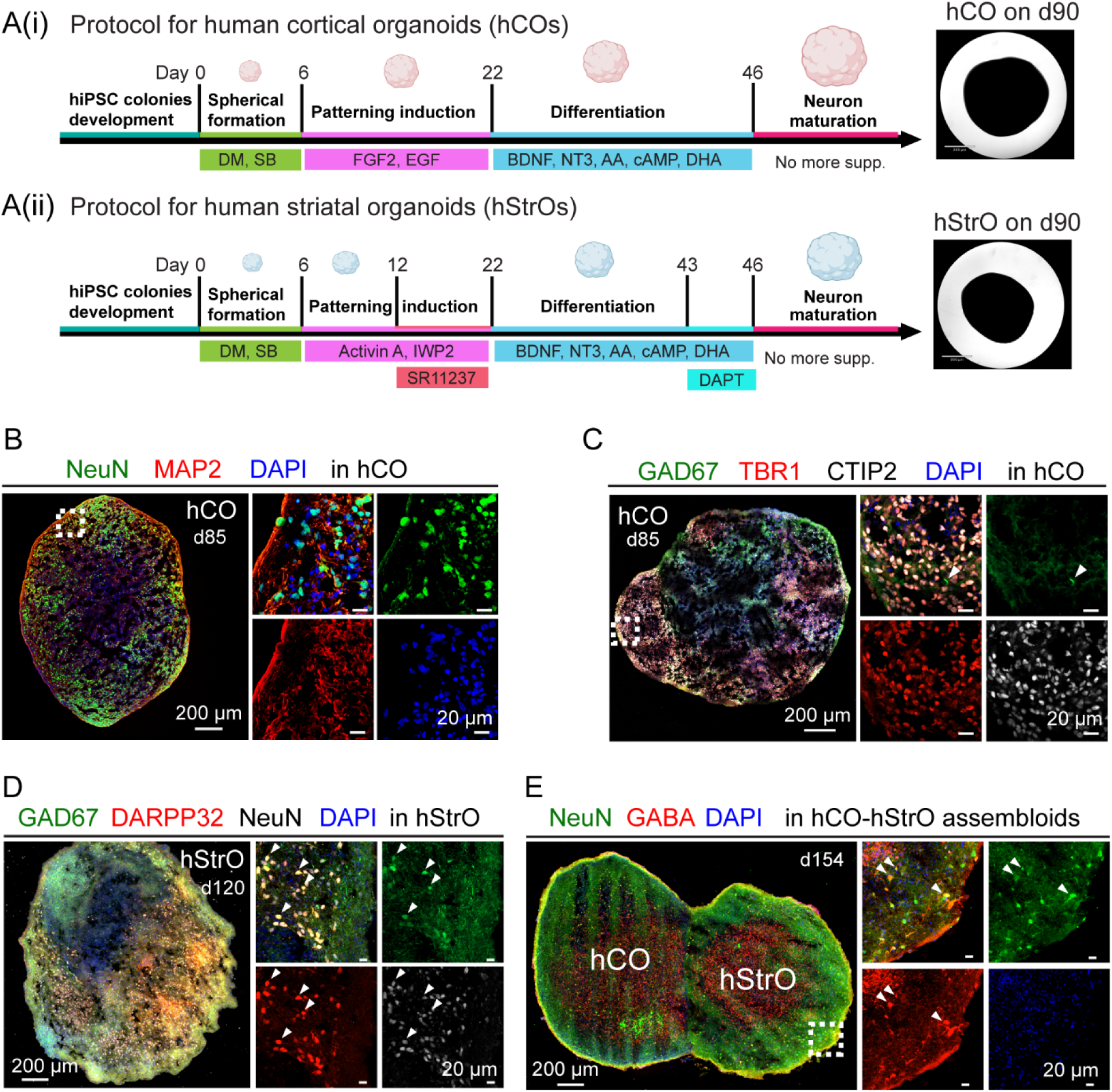
Generation and characterization of human cortical and striatal organoids. Related to. Figure 1 (A(i)) Protocol for human cortical organoids (hCOs). Schematic timeline depicting hCO differentiation: human induced pluripotent stem cells (hiPSCs) undergo spheroid formation (day 6), patterning induction (day 22), differentiation (day 46), and neuronal maturation. Key supplements include dual SMAD inhibitors (DM/SB), fibroblast growth factor 2 (FGF2), epidermal growth factor (EGF), brain-derived neurotrophic factor (BDNF), neurotrophin-3 (NT3), ascorbic acid (AA), cyclic adenosine monophosphate (cAMP), and docosahexaenoic acid (DHA). Right panel: Brightfield image of an hCO, scale bar: 860 µm. (Aii) Protocol for human striatal organoids (hStrOs). Similar differentiation timeline with additional factors, including Activin A, inhibitor of Wnt production-2 (IWP2), and SR11237 for patterning induction, followed by the γ-secretase inhibitor DAPT for late-stage differentiation. Right panel: Brightfield image of an hStrO, scale bar: 860 µm. (B) Immunostaining for neuronal markers in hCO on day 85 showing neuronal nuclei (NeuN, green, a marker for mature neurons), microtubule-associated protein 2 (MAP2, red, a dendritic marker), and 4′,6-diamidino-2-phenylindole (DAPI, blue, a nuclear marker). High-magnification images show individual marker localization within the spheroid. (C) Neuronal type characterization in hCO. Immunostaining reveals sparse expression of glutamate decarboxylase 67 (GAD67, green, a marker for GABAergic neurons) and strong expression of T-box brain transcription factor 1 (TBR1, red, a marker for deep-layer cortical neurons), COUP-TF interacting protein 2 (CTIP2, white, a marker for subcortical projection neurons), and DAPI (blue). High-magnification images illustrate distinct neuronal populations within the spheroid. (D) Immunostaining of hStrO on day 120 showing GAD67 (green), dopamine- and cAMP-regulated phosphoprotein 32 (DARPP32, red, a marker for striatal projection neurons), NeuN (white), and DAPI (blue). High-magnification images highlight the spatial distribution of these neuronal markers. (E) Immunostaining of hCO-hStrO assembloids, showing NeuN (green), gamma-aminobutyric acid (GABA, red, a marker for GABAergic neurons), and DAPI (blue). The cortical and striatal regions are labeled as hCO and hStrO, respectively. High-magnification images display marker localization at the fusion site, indicating the integration of cortical and striatal neuronal populations.

**Figure S2.**
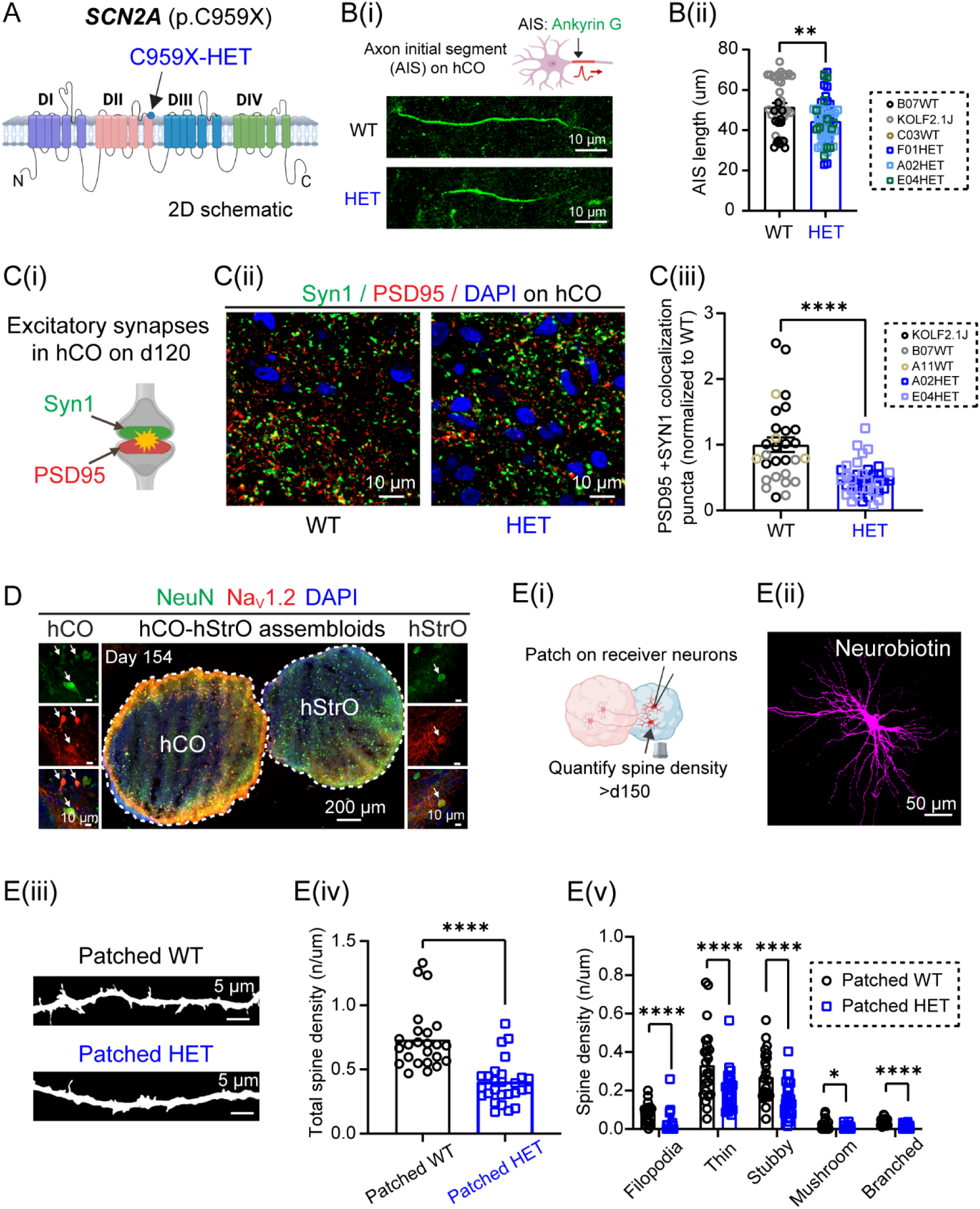
*SCN2A* deficiency disrupts AIS length, synaptic connectivity, and spine morphology in hCO-hStrO assembloids. Related to Figure 2. (A) Schematic representation of *SCN2A* highlighting the p.C959X nonsense mutation site. (B) AIS morphology. (B(i)) Schematic and representative images of AISs labeled by Ankyrin-G (green) in WT and HET *SCN2A-C959X* neurons. (B(ii)) AIS length quantification (WT: n = 42 neurons from B07, C03, KOLF2.1J hiPSC lines; HET: n = 56 neurons from F01, A02, E04 hiPSC lines). Unpaired Welch’s t test, **p < 0.01. (C) Impaired excitatory synapse formation in *SCN2A*-mutant hCOs. (C(i)) Schematic of excitatory synapses labeled by Syn1 (green, presynaptic) and PSD95 (red, postsynaptic). (C(ii)) Representative images of excitatory synapses (PSD95^+^/Syn1^+^ colocalization) in WT and HET *SCN2A-C959X* hCOs. Scale bar: 10 µm. (C(iii)) Quantification of excitatory synapses. WT: 29 images from 9 organoids (KOLF2.1J, B07, A11); HET: 37 images from 10 organoids (A02, E04). Unpaired t-test, ***p < 0.001. (D) Immunostaining of NeuN (green), Na_V_1.2 (red), and DAPI (blue) in hCO-hStrO assembloids, showing colocalization of Na_V_1.2 with NeuN in both hCO and hStrO, indicating Na_V_1.2 expression in neuronal cell bodies. (E) Reduced dendritic spine density in *SCN2A*-mutant striatal neurons in hCO-hStrO assembloid slices. (E(i)) Schematic of patch-clamp recording on hStrO receiver neurons and dendritic spine density quantification. (E(ii)) Representative images of patched neurons labeled with neurobiotin (magenta). (E(iii)) Representative images of dendritic spines from patched neurons. (E(iv)) Quantification of total receiver spine density. WT (gray, n = 24 dendrites, from 4 assembloids, A11, KOLF2.1J); HET *SCN2A-C959X* (blue, n = 27 dendrites, from 5 assembloids, A02, E04). Mann-Whitney test: ****pc<c0.0001. (E(v)) Spine subtype analysis (filopodia, thin, stubby, mushroom, branched). Multiple Mann-Whitney test, *p < 0.05, ****p < 0.0001. Data are represented as mean ± SEM.

**Figure S3.**
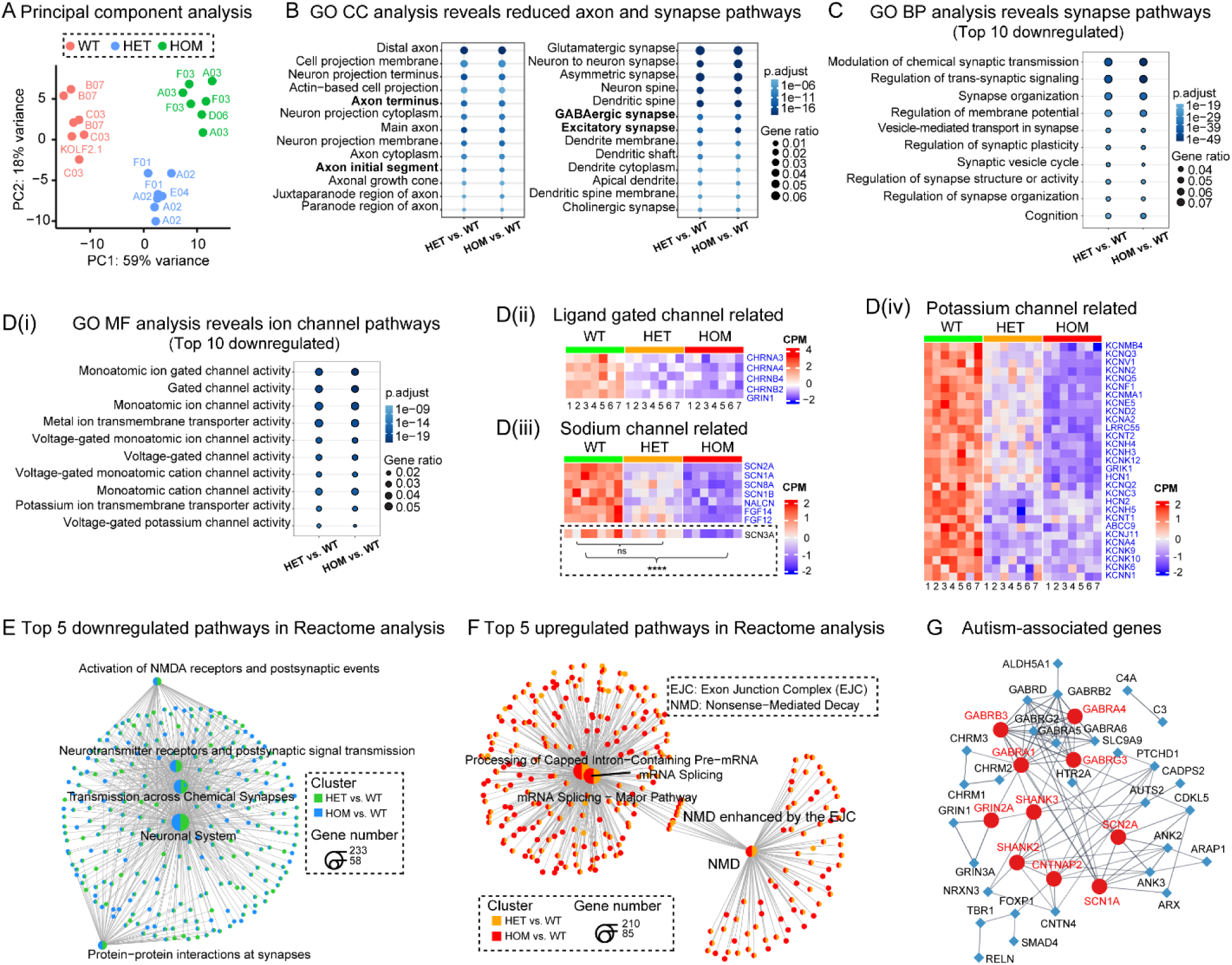
Dysregulated gene expressions and pathway alterations in *SCN2A*-deficient assembloids. Related to Figure 3. (A) Global transcriptomic alterations. Principal component analysis (PCA) of transcriptomic data showing clustering of WT (KOLF2.1J, B07, C03 lines), HET (A02, E04, F01 lines), and HOM (A03, D06, F03 lines) assembloids based on global gene expression. Each dot represents a sample (n = 7 assembloids/group). (B) Gene ontology (GO) cellular component (CC) analysis of overlapping downregulated genes, revealing reductions in axon terminal and axon initial segments; excitatory synapses, and GABAergic synapses. (C) Disrupted synapse-related pathways in *SCN2A-C959X* assembloids. Top 10 overlapping downregulated pathways in GO biological processes (BP) in HET vs. WT and HOM vs. WT comparisons. (D) Disrupted ion channel-related pathways in *SCN2A-C959X* assembloids. (D(i)) The top 10 overlapping downregulated pathways in GO molecular functions (MF) reveal ion channel gene dysregulation in HET and HOM assembloids. Heatmaps of overlapping downregulated ion channel genes (blue), categorized into ligand-gated channel-related (D(ii)), voltage-gated sodium channel-related (D(iii)), and voltage-gated potassium channel-related (D(iv). Highlighted *SCN3A* showed no significant changes in the HET group but was significantly reduced in the HOM group. (E) Synaptic dysfunction in HET and HOM assembloids. Network plot of the top 5 Reactome pathways enriched in downregulated DEGs, primarily affecting NMDA receptor activity, neurotransmitter signaling, synaptic transmission, and neuronal interactions, suggesting disrupted synaptic function in *SCN2A*-deficient assembloids. (F) Increased RNA surveillance in *SCN2A-C959X* assembloids. Network plot of the top 5 Reactome pathways enriched in upregulated DEGs, revealing associations with mRNA splicing, nonsense-mediated decay (NMD), and exon junction complex (EJC) function. (G) Network analysis of ASD-associated genes, depicting interactions between differentially expressed ASD-related genes in *SCN2A*-mutant assembloids. Top ten hub genes are highlighted in red.

## EXPERIMENTAL MODEL AND SUBJECT DETAILS

### The hiPSC Lines

*SCN2A* c.2877C>A (p.Cys959Ter) mutant hiPSC lines were generated using CRISPR/Cas9-mediated genome editing^34,35^ in early passage (p2) KOLF2.1J reference iPSCs^36^. We conducted recharacterization of pluripotency assays and genome integrity to ensure the quality of hiPSC^37^. This study utilized four isogenic WT control lines (KOLF2.1J, C03, B07, and A11), three heterozygous *SCN2A* mutant lines (A02, E04, and F01), and three homozygous *SCN2A* mutant lines (A03, D06, and F03). The successful introduction of the mutation was confirmed by Sanger sequencing and genotyping for each batch of hiPSC-derived organoids or assembloids.

### Antibodies

For immunostaining, primary antibodies used include Anti-NeuN (Chicken, GeneTex, GTX00837, 1:1000; Rabbit, Cell Signaling, 24307S, 1:1000), Anti-TBR1 (Rabbit, Abcam, ab31940, 1:300), Anti-MAP2 (Chicken, Novus Biologicals, NB300-213, 1:1000; Mouse, Millipore, MAB378, 1:200), Anti-GABA (Rabbit, Sigma-Aldrich, A2052, 1:1000), Anti-GAD67 (Mouse, Sigma-Aldrich, MAB5406-25UG, 1:100), Anti-Ctip2 (Rat, Abcam, ab18465, 1:300), Anti-DARPP32 (Rabbit, Abcam, ab40801, 1:200), Anti-Ankyrin-G (Mouse, Antibodies Inc., 75-146-020, 1:200), Anti-SCN2A (Rabbit, Sigma-Aldrich, ZRB1300, 1:200), HA-Tag (Rabbit, Cell Signaling, 3724T, 1:200), Synapsin 1 (Mouse, Synaptic Systems, 106011, 1:800), PSD-95 (Rabbit, Thermo Fisher, 51-6900, 1:800). Secondary antibodies included Goat anti-Chicken Alexa Fluor 647 (Thermo Fisher, A32933, 1:500), Goat anti-Rabbit Alexa Fluor 647 (A21244, 1:500), Goat anti-Mouse Alexa Fluor 647 (A21235, 1:500), Goat anti-Rat Alexa Fluor 555 (A21434, 1:500), Goat anti-Rabbit Alexa Fluor 488 (A11034, 1:500), Goat anti-Rabbit Alexa Fluor Plus 555 (A32732, 1:500), Goat anti-Mouse Alexa Fluor Plus 555 (A32727, 1:500), and Goat anti-Mouse Alexa Fluor 488 (A11001, 1:500).

For Western blotting, primary antibodies included Rabbit anti-SCN2A (Na_V_1.2) (Alomone Labs, ASC-002, 1:200), Rabbit anti-*SCN2A* (Sigma-Aldrich, ZRB1300, 1:200), and Mouse anti-β-Actin (Thermo Fisher, MA5-15739, 1:1000). Secondary antibodies used were IRDye® 800CW Goat anti-Rabbit IgG (LI-COR, 926-32211, 1:2500) and IRDye® 800CW Goat anti-Mouse IgG (926-32210, 1:5000).

## METHOD DETAILS

### Genotyping

hiPSCs, organoids, and assembloids were collected and genotyped. DNA was extracted using a tissue DNA extraction kit (Macherey-Nagel, Bethlehem, USA). Gene-specific PCR was performed using the following primers: Forward (5’–3’): TTGAGACAGTTACCTGTACATTTGC; Reverse (5’–3’): TAATAGACAATAGGAAGTGGCCTTG. PCR reactions (25 µL) were prepared with PCR Master Mix, primers, template DNA, and nuclease-free water, followed by amplification under the thermal cycling conditions: 95°C for 2 minutes, then 35 cycles of 95°C for 30 seconds, 55°C for 30 seconds, and 72°C for 1 minute, with a final extension at 72°C for 10 minutes, and hold at 4°C. The 700-bp PCR product was subjected to Sanger sequencing, confirming genome editing: WT retained cytosine (C) at the mutation site, heterozygous lines displayed overlapping peaks, and homozygous lines showed a complete C-to-A substitution, introducing a premature stop codon.

### Generation of hStrOs and hCOs from hiPSCs

#### Generation of 3D neural organoids

Feeder-free hiPSCs colonies were seeded on Matrigel (Corning, # 354230) in StemFlex medium (Thermo Fisher, # A3349401) with daily media changes. Cells were passaged every 4–

5 days using Versene solution (Thermo Fisher, #15040066). Quality control assessments included Sanger sequencing, karyotyping, and immunocytochemistry. Undifferentiated hiPSC colonies displayed normal and homogenous morphology with defined edges and minimal spontaneous differentiation.

Organoids were generated following a modified Muria protocol^15^. hiPSCs were dissociated into single cells using Accutase (Thermo Fisher, # NC9839010) and seeded at 10,000 cells/well in ultralow-attachment 96-well plates (Corning, #CLS7007) with Essential 8 medium (Thermo Fisher, #A1517001) supplemented with 10 μM ROCK inhibitor Y27632 (Selleck Chemicals, #S1049). Plates were centrifuged at 100 g for 3 minutes and incubated at 37°C with 5% CO_2_. After 24 hours, organoids were transferred to Essential 6 medium (Thermo Fisher, #A1516401) supplemented with 2.5 μM dorsomorphin (Sigma-Aldrich, #P5499), 10 μM SB-431542 (R&D Systems, #1614), and 1.25 μM XAV-939 (Tocris, #3748) for 5 days with daily media changes.

#### Patterning and differentiation into hCO and hStrOs

On day 6, organoids were transferred into ultralow-attachment 6-well plates (Corning, # 3471) with neural medium containing Neurobasal-A (Thermo Fisher, #10888022), B-27 without vitamin A (Thermo Fisher, #12587010), GlutaMAX (1:100, Thermo Fisher, #35050061), and penicillin-streptomycin (1:100, Thermo Fisher, #15140122). For hCOs, this medium was supplemented with 20 ng/mL FGF2 (R&D Systems, # 233-FB-500) and 20 ng/mL EGF (R&D Systems, # 236-EG). For hStrOs, this medium was supplemented with 2.5 μM WNT inhibitor IWP-2 (Selleck Chemicals, #S7085) and 50 ng/mL recombinant Activin A (PeproTech, #120-14P). On day 12, 100 nM SR11237 (Tocris, #3411) was added to hStrOs.

#### Neuronal differentiation and maturation

From day 22, neural progenitors were differentiated into neurons for both hCO and hStrO by supplementing the neural medium with BDNF (20 ng/mL, PeproTech, #450-02), NT-3 (20 ng/mL, PeproTech, #450-03), ascorbic acid (200 μM, Wako, #323-44822), Dibutyryl-cAMP (50 μM, Santa Cruz, #sc-201567A), and DHA (10 μM, MilliporeSigma, #D2534). From days 42– 45, 2.5 μM DAPT (Stemcell Technologies, #72082) was added to hStrO along with BDNF, NT-3, ascorbic acid, cAMP, and DHA. From day 46 onward, cultures were maintained in a neural medium containing B-27 Plus Supplement (Thermo Fisher, #A3582801) with medium changes every 4–5 days.

### Viral Labeling and Live-Cell Imaging of Cortico-Striatal Assembloids

#### Generation of cortico-striatal assembloids^13^

To generate cortico-striatal assembloid, one hCO and one hStrO were transferred into a 1.5-mL Eppendorf tube containing 1 mL NM medium following viral labeling, incubated at 37°C for 3–4 days with a complete medium change on day 2. Once formed, assembloids were transferred to ultralow-attachment plates using a P1000 pipette with a cut tip to accommodate their size. Assembly was conducted between days 60–76 of differentiation.

#### Anterograde and retrograde tracing

For anterograde labeling, on day 60, hCOs were labeled with AAV1-hSyn-Cre (Addgene #105553), while hStrOs were labeled with AAV1-Ef1a-DIO-mScarlet (Addgene #131002) and AAV9-hSyn-EGFP (Addgene #50465). For retrograde labeling, hStrOs were labeled with AAV1-hSyn-Cre, while hCOs were labeled with AAV1-Ef1a-DIO-mScarlet and AAV9-hSyn-EGFP. Assembloids were maintained in culture with medium changes every 4 days. After 1 month, labeled assembloids were transferred to glass-bottom 24-well plates (Cellvis, #P24-0-N) in BrainPhys medium (STEMCELL, #5790) and imaged using a Zeiss LSM 900 confocal microscope (10× objective, Z-stack scanning) following a 15-minute equilibration at 37°C and 5% CO_2_.

#### Axon projection imaging

To assess projections from hCO to hStrO, hCOs were labeled with AAV1-hSyn-mScarlet (Addgene #131001) on day 60, assembled with hStrO on day 65, and imaged on days 10, 20, 30, and 42 post-fusion using Zeiss LSM 900 (10× objective, 0–500 μm depth, Z-stack scanning). Axonal projections were quantified by measuring the percentage of mScarlet coverage in hStrO using Fiji (https://fiji.sc/).

#### Receiver and non-receiver neuron labeling

hCOs were labeled with AAV1-hSyn-Cre, while hStrOs were labeled with AAV1-Ef1a-DIO-mScarlet and AAV9-hSyn-EGFP on day 60, followed by fusion on day 65. After 1 month, dendritic spine imaging was conducted using Zeiss LSM 900 (20× objective, 6× zoom, Z-stack scanning) following 15 minutes of equilibration at 37°C and 5% CO_2_. EGFP^+^/mScarlet*^−^* neurons were classified as non-receivers, while mScarlet^+^ neurons were designated as receivers. At least three dendrites per assembloid were analyzed for spine density and morphology using Neurolucida 360 and Neurolucida Explorer (MBF Bioscience).

#### Dendritic spine imaging in receiver neurons

hCOs were labeled with AAV1-hSyn-Cre, while hStrOs were labeled with AAV1-Ef1a-DIO-mScarlet on day 60, followed by fusion on day 65. After 1 month, assembloids were transferred to BrainPhys medium, and receiver (mScarlet^+^) neurons were imaged at 0–500 μm depth. Dendritic spines were visualized with a 20× objective and 6× zoom, and neuronal complexity, spine density, and morphology were assessed by Sholl analysis using Neurolucida 360 and Neurolucida Explorer (MBF Bioscience).

### Patch-Clamp Recordings

#### Acute slice preparations

Slices were prepared from 3–6-month-old brain organoids and hCO-hStrO assembloids. Organoids and assembloids were rapidly embedded in 4% agarose in slicing solution containing (in mM): 110 choline chloride, 2.5 KCl, 1.25 NaH_2_PO_4_, 25 NaHCO_3_, 0.5 CaCl_2_, 7 MgCl_2_, 25 glucose, 1 sodium ascorbate and 3.1 sodium pyruvate (pH 7.4, 305–315 mOsm, bubbled with 95% O_2_ and 5% CO_2_). Coronal slices (200 μm thick) were prepared using a vibratome (Leica VT1200 S, Germany). Slices were incubated for 10 minutes at 33°C in slicing solution, then transferred to artificial cerebrospinal fluid (aCSF; in mM; 125 NaCl, 2.5 KCl, 1.25 NaH_2_PO_4_, 25 NaHCO_3_, 2.0 CaCl_2_, 2.0 MgCl_2_, 10 glucose; pH 7.4, 305–315 mOsm, bubbled with 95% O_2_ and 5% CO_2_) for 10–20 minutes at 33°C before storage at room temperature for at least 30 minutes prior to recording.

#### Whole-cell electrophysiology in assembloid slices

Slices were placed in a recording chamber continuously perfused with aCSF at 32–33°C (2–3 mL/min). Neurons were visualized with an IR-DIC microscope (Olympus BX-51WI) equipped with an IR-2000 camera (Dage-MTI). Somatic whole-cell patch-clamp recordings were performed on striatal medium spiny neurons (MSNs) and cortical pyramidal neurons. MSNs were identified by their characteristic morphology, including medium-sized, polygonal/dendritic spiny somata, while cortical pyramidal neurons were distinguished by their prominent apical dendrite. Thin-wall borosilicate pipettes (BF150-110-10, Sutter Instruments) with an open-tip resistance of 3–5 MΩ were fabricated using a P-1000 puller (Sutter Instruments).

Voltage-clamp recordings of spontaneous excitatory postsynaptic currents (sEPSCs) were performed to evaluate the synaptic activity. The internal solution consisted of (in mM): 120 CsMeSO_3_, 4 MgCl_2_, 0.2 EGTA, 10 HEPES, 4 Na_2_ATP, 0.3 Tris_3_-GTP, 14 Tris_2_-phosphocreatine, adjusted to pH 7.25 with CsOH (295–305 mOsm). sEPSCs were recorded for 3 minutes in gap-free mode with membrane potentials held at *V*_hold_ = −80 mV.

Recordings were performed using an Axon MultiClamp 700B amplifier (Molecular Devices), and data were acquired with pClamp 11.4 software, filtered at 2 kHz, and sampled at 33 kHz with an Axon Digidata 1550B plus HumSilencer digitizer (Molecular Devices). Series resistance (Rs) was maintained at 15–30 MΩ, and recordings with unstable Rs (>20%) were excluded. Data files were saved in ABF 1.8 format and analyzed using the Mini Analysis Program (v6.08, Synaptosoft).

For chemogenetics experiments, hCOs were labeled with AAV9-hSyn-hM3D(Gq)- mCherry (Addgene #50474), while hStrOs were labeled with AAV9-hSyn-EGFP on day 60, followed by assembly on day 65. After two months, assembloids were prepared for sEPSC recordings. EGFP⁺ MSNs in hStrO, surrounded by mCherry-labeled axons, were selected for recording. Baseline activity was recorded for 3 minutes, followed by a bath application of 10 µM CNO for 3 minutes, and a final 3-minute washout.

The internal solution for cell labeling included 0.1–0.2% neurobiotin. After recordings (∼30 minutes), slices were fixed in 4% paraformaldehyde (pH 7.4) for 20–30 minutes at room temperature, washed in PBS, and incubated overnight at 4°C with Alexa 647-conjugated streptavidin (1:250 in blocking solution)^38^. Neuronal morphology, including dendritic spines, of patched neurons, was imaged under Zeiss LSM 900.

#### In vitro whole-cell electrophysiology in dissociated neurons

Brain organoids were dissociated into single cells using a modified Worthington Papain Dissociation Kit (Papain Dissociation System, Worthington, LK003150). 3∼5 randomly selected mature hCOs (>110 days old) were transferred to 60 mm dishes, and the excess medium was aspirated before adding 5 mL of Papain-DNase solution. The tissue was minced into <1 mm pieces and incubated at 37°C, 5% CO_2_, shaking at 80 rpm for 30 minutes, and then gently triturated, followed by an additional 10-minute incubation. The resulting suspension was transferred to a 15 mL conical tube, allowing debris to settle, and the supernatant was mixed with Inhibitor solution before centrifugation at 300g for 7 minutes. The pellet was resuspended in Neurobasal medium, filtered through a 40 μm strainer (CELLTREAT Scientific Products, #229481), and counted. Approximately 1–2 × 10^5^ cells were seeded onto 12-mm round coverslips (Neuvitro, #GG-12-1.5-Pre) in 24-well plates. Neurons were cultured in a Neurobasal medium with a B-27 supplement (without vitamin A) for 7 days, with half-medium changes every other day. From day 7, cells were maintained in the Neurobasal medium with B-27 Plus Supplement, with half-medium changes twice a week.

For whole-cell current-clamp recordings, the internal solution contained (in mM): 122 KMeSO_4_, 4 KCl, 2 MgCl_2_, 0.2 EGTA, 10 HEPES, 4 Na_2_ATP, 0.3 Tris-GTP, and 14 Tris-phosphocreatine, adjusted to pH 7.25 with KOH, 295–305 mOsm. Recordings were conducted using an Axon MultiClamp 700B amplifier (Molecular Devices), acquired at 50 kHz, and filtered at 2 kHz with pClamp 11.4 software. The sag ratio, input resistance, and firing properties were assessed using a series of 400-ms hyperpolarizing and depolarizing current steps from −40 pA to +140 pA in 20-pA increments, with 5-s sweeps at either the normal resting membrane potential (RMP) or −70 mV.

Input resistance (*R*_input_) was calculated as:

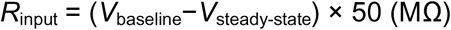

where *V*_baseline_ is the RMP or −70 mV, and *V*_steady-state_ is the voltage recorded before the end of the −20-pA stimulus.

### High-Density Multielectrode Array (HD-MEA) Recording of Cortico-Striatal Assembloids

#### HD-MEA preparation and assembloid seeding

HD-MEAs (3Brain, 6-well plates) were cleaned with 1% Tergazyme (200 μL/well, 37°C, 1 hour), rinsed three times with ultrapure water, disinfected in 70% ethanol for 1 hour, dried, and incubated overnight in PBS at room temperature. HD-MEAs were coated sequentially with poly-L-ornithine (50 μg/mL; Sigma-Aldrich, #P4957) overnight, washed, and further coated with laminin (50 μg/mL, ≥2 hours at 37°C).

Cortico-striatal (hCO-hStrO) assembloids (∼4 months old) were transferred onto HD-MEAs using a wide-bore pipette and allowed to settle for 2 hours. To promote attachment, 20 μL of medium was added every 2 hours for 8 hours, followed by 2 mL of medium the next day. Cultures were maintained in BrainPhys medium supplemented with N2, B27 Plus, 50 μM cAMP, and 200 μM ascorbic acid, with biweekly media changes. Recordings were conducted after 2 weeks of culture.

#### Electrophysiological recordings and analysis

On the day of recording, the media was refreshed, and the plate was incubated at 37°C with 5% CO_2_ for 15 minutes. Extracellular activity was recorded for 5 minutes using Brainwave V software (v5.6, 3Brain Switzerland) in the HyperCAM Alpha multi-well system. Recordings were acquired from 2304 × 2304 electrode HD-MEAs (60 μm pitch, 2.9 × 2.9 mm^2^ recording area) at a 10 kHz sampling rate, with 100 Hz high-pass and a 20–5000 Hz band-pass filtering. Fast Fourier Transform (FFT) with a Hamming window was applied for spectral analysis.

Spikes were detected using an 8.0 standard deviation threshold, with a peak lifetime of 2.0 ms, refractory period of 1.0 ms, and pre-peak wave duration of 1.0 ms. Electrodes with spike frequencies < 0.083 Hz (5 spikes/min) were excluded. Bursts were identified as ≥5 spikes with an interspike interval ≤100 ms, while network bursts were detected via a recruitment-based algorithm requiring ≥10% electrode activation and a minimum burst size of 50 spikes. Spike sorting was performed using Principal Component Analysis (PCA, 3 components) and K-Means clustering with Gap Statistics.

### Cryoprotection, Immunocytochemistry, and Imaging Analysis

#### Sample preparation

Brain organoids and assembloids were fixed in 4% paraformaldehyde (PFA) in PBS overnight at 4°C, washed in PBS, and transferred to 30% sucrose-PBS for 2–3 days until fully submerged. Samples were then equilibrated in a 1:1 mixture of optimal cutting temperature (OCT) compound (Tissue-Tek, 4583, Sakura Finetek) and 30% sucrose-PBS before embedding. After embedding, cryosections (20–40 μm thick) were obtained using a Leica CM1850 cryostat. Slices on glass coverslips for the 2D culture were fixed in 4% PFA for 20 minutes at room temperature.

#### Immunostaining^39^

Cryosections were washed (3×, 5 minutes) in PBS, permeabilized, and blocked in either 0.5% Triton X-100 and 5% normal goat serum in PBS or 4% Block-Ace (Dainippon Sumitomo Pharma, UK-B80) with 0.05% Tween-20 in PBS for 1 hour at room temperature. Primary antibodies were applied overnight at 4°C, followed by PBS washes (3×, 10 minutes). Samples were then incubated with Alexa Fluor-conjugated secondary antibodies for 1 hour at room temperature in a blocking buffer. Sections were mounted with DAPI-containing Antifade Mounting Medium (VECTASHIELD, H-2000) and sealed with glass coverslips. Images were acquired using an LSM900 confocal fluorescence microscope equipped with an air scan module (Carl Zeiss, Jena, Germany).

#### Axon initial segment (AIS) length quantification^23^

Images were captured using a 63× oil-immersion objective, with Z-stacks collected from at least three regions per organoid. AIS length was measured from maximum intensity projections of Z-stacks spanning the entire neuron, based on ankyrin-G immunofluorescence. Only AIS structures with clearly defined start and end points were analyzed using the segmented line tool in Fiji.

#### Synaptic density quantification

Synaptic density was analyzed using a 63× oil-immersion objective with 1.3× zoom with an air scan module, using identical imaging parameters for both WT and mutant. For each organoid, 3–6 regions of interest were captured. Colocalization of excitatory presynaptic (SYN1) and postsynaptic (PSD95) puncta was assessed using built-in analysis software (Nikon system), with consistent analysis settings applied for both WT and mutant every batch. Synaptic density of colocalized puncta was normalized to the WT group to evaluate changes in the mutant group.

### Western Blotting

Organoids and assembloids were homogenized in ice-cold RIPA buffer (Thermo Fisher, 89901) with protease and phosphatase inhibitors (Thermo Fisher, A32953) and centrifuged at 10,000× g for 20 minutes at 4°C. Protein concentrations were determined using the Pierce^TM^ BCA Protein Assay Kit (Thermo Fisher, 23225). Proteins were denatured in 1× Laemmli buffer (62.5 mM Tris-HCl (pH 6.8), 2% SDS, 5% glycerol, 0.05% bromophenol blue) by boiling at 95°C for 5 minutes. Then 50 μg of per sample was loaded on 8% SDS-PAGE gels at 80–120 V and transferred onto PVDF membranes (Thermo Fisher, PB9220) at 300 mA for 2.5 hours at 4°C. Membranes were blocked with 5% nonfat milk in TBST (Tris-buffered saline with 0.1% Tween 20) for 1 hour at room temperature and incubated overnight at 4°C with primary antibodies diluted in Intercept^®^ T20 (TBS) Antibody Diluent (LI-COR Biosciences, 927-65001). The next day, blots were washed (3×, 10 minutes) in TBST, incubated with secondary antibodies for 1 hour at room temperature, and washed again (3×, 10 minutes). Immunoreactive bands were visualized using the Odyssey^®^ CLx Imaging System (LI-COR Biosciences) and analyzed with Fiji. Protein levels were normalized to β-actin and further compared.

### RNA Isolation, Reverse Transcription, and qPCR Analysis

Total RNA was extracted from hCO-hStrO assembloids using the RNeasy Mini Kit (QIAGEN, #74104) following the protocol by the manufacturer. RNA integrity and concentration were assessed using a NanoDrop spectrophotometer (Thermo Fisher). Reverse transcription was performed using the Maxima First Strand cDNA Synthesis Kit (Thermo Fisher, #K1672) under optimal conditions to ensure efficient cDNA synthesis.

Quantitative PCR (qPCR) was conducted using PowerUp SYBR Green Master Mix (Thermo Fisher, #A25777) and gene-specific primers on a C1000 Touch PCR thermal cycler (Bio-Rad), following the recommended protocol by the manufacturer. Thermal cycling conditions were as follows: initial denaturation at 95°C for 1 minute, followed by 45 cycles of 95°C for 15 seconds and 60°C for 60 seconds. For *SCN2A* amplification included internal *SCN2A* primers (Forward: GAGACCATGTGGGACTGTATG; Reverse: AAGGCCAAGAAGAGGTTCAG), codon-optimized *SCN2A* primers (Forward: GTGTTTTGCCTCTCCGTGTT; Reverse: ATTTCCGTCCAGGGAGTTGT), and total *SCN2A* primers (Forward: GGATACATCTGTGTGAAGGC; Reverse: CTGTTCCTCATAGGCCAT).

Gene expression levels were normalized to GAPDH mRNA, as an endogenous control, calculated as (ΔCt):

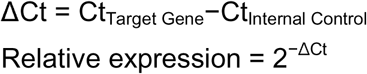

All reactions were performed duplicated to ensure reproducibility, and data were analyzed using Bio-Rad CFX Manager software.

### Bulk RNA Sequencing and Analysis

#### RNA extraction and library preparation

RNA was extracted from 5-month-old hCO-hStrO assembloids derived from three wild-type (WT) lines (KOLF2.1J, B07, C03), three heterozygous (HET) *SCN2A-C959X* lines (A02, E04, F01), and three homozygous (HOM) *SCN2A-C959X* lines (A03, D06, F03), totaling 21 assembloids (7 per genotype) from two independent batches. Polyadenylated (Poly(A)^+^) RNA was isolated from 100–250 ng of total RNA using the NEBNext^®^ Poly(A) mRNA Magnetic Isolation Module (New England Biolabs). RNA fragmentation and elution were performed directly from the oligo dT beads as part of the library construction process using the xGen RNA Library Preparation Kit (Integrated DNA Technologies, IDT) according to the manufacturer’s instructions. Prepared libraries were pooled and sequenced on an Illumina NovaSeq X+ system using a 300-cycle 25B flow cell, generating 30–36 million paired-end reads per sample. FASTQ files were generated using BCL2FASTQ software (version 1.8.4) for downstream analysis.

#### Preprocessing, alignment, and genotype verification

Raw FASTQ files were processed using fastp (v0.23.2)^40^ to remove adapter sequences and trim low-quality bases (Phred score <30). Reads shorter than 50 bp after trimming were discarded. The remaining reads were aligned to the GRCh38 human reference genome (Ensembl release 104) using the STAR Aligner (v2.7.10a)^41^ in two-pass mode to improve splice junction detection. To confirm the genotype of the C959X mutation in *SCN2A*, variant calling was performed using GATK HaplotyperCaller (v4.2.2.0)^42^ with Joint Genotyping^43^. This analysis ensured the accurate classification of samples into HOM, HET, and WT groups.

#### Gene quantification and normalization

Read assignment to genomic features was performed using featureCounts (v1.6.1)^44^ in paired-end, reverse-stranded mode. Initial exploratory analysis was conducted with DESeq2

(v1.34.0)^45^ in R (v4.1.3)^46^, to evaluate sample clustering and overall consistency. The raw counts’ matrix was filtered to retain genes with at least 5 reads in two samples for batch effect correction in RUVSeq (v1.28.0)^47^ by computing deviance residuals. Data normalization was performed using the upper-quartile method via the betweenLaneNormalization function.

#### Differential expression and pathway analysis

A generalized linear model (GLM) regression approach was applied to the count data, incorporating mutation genotype as a covariate while adjusting for batch effects using RUVSeq factors (k = 5). edgeR (v3.36.0)^48^ was used to fit a quasi-likelihood negative binomial model for differential expression analysis. Statistical significance was determined using the Benjamini-Hochberg method for multiple hypothesis correction. Genes with an FDR < 0.05 were classified as differentially expressed (DEs). DE genes were analyzed for enrichment in KEGG and Reactome pathways, as well as Gene Ontology (GO) terms, using clusterProfiler (v4.10.0)^49^ in R (v4.3.2). The background gene set included all genes detected after RUVSeq correction. Ingenuity Pathway Analysis (IPA)^50^ was performed to provide curated insights into biological pathways and disease associations.

#### Data visualization

Heatmaps: Batch-effect corrected counts per million (CPM) values were extracted using edgeR and visualized with the ComplexHeatmap package (v2.14.0)^51^ in R (v4.2.1).

Volcano Plots: Differential expression results were visualized using the EnhancedVolcano package (v1.16.0)^52^. Network and Dot Plots: Pathway enrichment results were displayed using Enrichplot (v1.22.0) and clusterprofiler (v4.10.0) to highlight key gene interactions. Hub genes among autism genes were identified and visualized using the cytoHubba (v0.1) module in Cytoscape (v3.10.2).

## QUANTIFICATION AND STATISTICAL ANALYSIS

Normality and variance similarity were measured by GraphPad Prism before the application of any parametric tests. For comparisons between two groups, either a two-tailed Student’s *t* test (parametric) or an unpaired two-tailed Mann-Whitney *U* test (non-parametric) was applied. For multiple comparisons, data were analyzed using one-way or two-way ANOVA with Tukey’s post-hoc correction (parametric) or Kruskal-Wallis test with Dunn’s multiple comparison correction (non-parametric), as appropriate. *Post hoc* tests were conducted only when the primary analysis showed statistical significance. Error bars in all figures represent the mean ± SEM. Statistical significance was defined as p < 0.05. Significance levels are denoted as follows: p < 0.05 is indicated as *, p < 0.01 is indicated as **, p < 0.001 is indicated as ***, and p < 0.0001 is indicated as ****.

## REFERENCES

1. Shaw, K.A. (2025). Prevalence and early identification of autism spectrum disorder among children aged 4 and 8 years—Autism and Developmental Disabilities Monitoring Network, 16 Sites, United States, 2022. MMWR. Surveillance Summaries 74.

2. Klei, L., Sanders, S.J., Murtha, M.T., Hus, V., Lowe, J.K., Willsey, A.J., Moreno-De-Luca, D., Yu, T.W., Fombonne, E., Geschwind, D., et al. (2012). Common genetic variants, acting additively, are a major source of risk for autism. Mol Autism 3, 9. 10.1186/2040-2392-3-9.

3. Sanders, S.J., Murtha, M.T., Gupta, A.R., Murdoch, J.D., Raubeson, M.J., Willsey, A.J., Ercan-Sencicek, A.G., DiLullo, N.M., Parikshak, N.N., Stein, J.L., et al. (2012). De novo mutations revealed by whole-exome sequencing are strongly associated with autism. Nature 485, 237–241. 10.1038/nature10945.

4. Satterstrom, F.K., Kosmicki, J.A., Wang, J., Breen, M.S., De Rubeis, S., An, J.Y., Peng, M., Collins, R., Grove, J., Klei, L., et al. (2020). Large-Scale Exome Sequencing Study Implicates Both Developmental and Functional Changes in the Neurobiology of Autism. Cell 180, 568–584.e523. 10.1016/j.cell.2019.12.036.

5. Uddin, M., Tammimies, K., Pellecchia, G., Alipanahi, B., Hu, P., Wang, Z., Pinto, D., Lau, L., Nalpathamkalam, T., Marshall, C.R., et al. (2014). Brain-expressed exons under purifying selection are enriched for de novo mutations in autism spectrum disorder. Nat Genet 46, 742–747. 10.1038/ng.2980.

6. Ben-Shalom, R., Keeshen, C.M., Berrios, K.N., An, J.Y., Sanders, S.J., and Bender, K.J. (2017). Opposing Effects on Na(V)1.2 Function Underlie Differences Between SCN2A Variants Observed in Individuals With Autism Spectrum Disorder or Infantile Seizures. Biol Psychiatry 82, 224–232. 10.1016/j.biopsych.2017.01.009.

7. Spratt, P.W.E., Ben-Shalom, R., Keeshen, C.M., Burke, K.J., Jr., Clarkson, R.L., Sanders, S.J., and Bender, K.J. (2019). The Autism-Associated Gene Scn2a Contributes to Dendritic Excitability and Synaptic Function in the Prefrontal Cortex. Neuron 103, 673–685 e675. 10.1016/j.neuron.2019.05.037.

8. Nelson, A.D., Catalfio, A.M., Gupta, J.P., Min, L., Caballero-Floran, R.N., Dean, K.P., Elvira, C.C., Derderian, K.D., Kyoung, H., Sahagun, A., et al. (2024). Physical and functional convergence of the autism risk genes Scn2a and Ank2 in neocortical pyramidal cell dendrites. Neuron 112, 1133–1149 e1136. 10.1016/j.neuron.2024.01.003.

9. Murugan, M., Jang, H.J., Park, M., Miller, E.M., Cox, J., Taliaferro, J.P., Parker, N.F., Bhave, V., Hur, H., Liang, Y., et al. (2017). Combined Social and Spatial Coding in a Descending Projection from the Prefrontal Cortex. Cell 171, 1663–1677 e1616. 10.1016/j.cell.2017.11.002.

10. Ma, Z.-H., Lu, B., Li, X., Mei, T., Guo, Y.-Q., Yang, L., Wang, H., Tang, X.-Z., Ji, Z.-Z., Liu, J.-R., et al. (2022). Atypicalities in the developmental trajectory of cortico-striatal functional connectivity in autism spectrum disorder. Autism 26, 1108–1122. 10.1177/13623613211041904.

11. Jourdon, A., Wu, F., Mariani, J., Capauto, D., Norton, S., Tomasini, L., Amiri, A., Suvakov, M., Schreiner, J.D., Jang, Y., et al. (2023). Modeling idiopathic autism in forebrain organoids reveals an imbalance of excitatory cortical neuron subtypes during early neurogenesis. Nat Neurosci 26, 1505–1515. 10.1038/s41593-023-01399-0.

12. Huang, W.K., Wong, S.Z.H., Pather, S.R., Nguyen, P.T.T., Zhang, F., Zhang, D.Y., Zhang, Z., Lu, L., Fang, W., Chen, L., et al. (2021). Generation of hypothalamic arcuate organoids from human induced pluripotent stem cells. Cell Stem Cell 28, 1657–1670 e1610. 10.1016/j.stem.2021.04.006.

13. Miura, Y., Li, M.Y., Birey, F., Ikeda, K., Revah, O., Thete, M.V., Park, J.Y., Puno, A., Lee, S.H., Porteus, M.H., and Pasca, S.P. (2020). Generation of human striatal organoids and cortico-striatal assembloids from human pluripotent stem cells. Nat Biotechnol 38, 1421–1430. 10.1038/s41587-020-00763-w.

14. Andersen, J., Revah, O., Miura, Y., Thom, N., Amin, N.D., Kelley, K.W., Singh, M., Chen, X., Thete, M.V., Walczak, E.M., et al. (2020). Generation of Functional Human 3D Cortico-Motor Assembloids. Cell 183, 1913–1929 e1926. 10.1016/j.cell.2020.11.017.

15. Miura, Y., Li, M.-Y., Revah, O., Yoon, S.-J., Narazaki, G., and Pacca, S.P. (2022). Engineering brain assembloids to interrogate human neural circuits. Nat Protoc 17, 15–35. 10.1038/s41596-021-00632-z.

16. Birey, F., Li, M.Y., Gordon, A., Thete, M.V., Valencia, A.M., Revah, O., Pasca, A.M., Geschwind, D.H., and Pasca, S.P. (2022). Dissecting the molecular basis of human interneuron migration in forebrain assembloids from Timothy syndrome. Cell Stem Cell 29, 248–264 e247. 10.1016/j.stem.2021.11.011.

17. Kim, J.I., Miura, Y., Li, M.Y., Revah, O., Selvaraj, S., Birey, F., Meng, X., Thete, M.V., Pavlov, S.D., Andersen, J., et al. (2024). Human assembloids reveal the consequences of CACNA1G gene variants in the thalamocortical pathway. Neuron 112, 4048–4059 e4047. 10.1016/j.neuron.2024.09.020.

18. Onesto, M.M., Kim, J.I., and Pasca, S.P. (2024). Assembloid models of cell-cell interaction to study tissue and disease biology. Cell Stem Cell 31, 1563–1573. 10.1016/j.stem.2024.09.017.

19. Steiner, H., and Tseng, K.Y. (2016). Handbook of basal ganglia structure and function (Academic Press).

20. Zhang, J., Chen, X., Eaton, M., Wu, J., Ma, Z., Lai, S., Park, A., Ahmad, T.S., Que, Z., Lee, J.H., et al. (2021). Severe deficiency of the voltage-gated sodium channel Na(V)1.2 elevates neuronal excitability in adult mice. Cell reports 36. 10.1016/j.celrep.2021.109495.

21. Harley, P., Kerins, C., Gatt, A., Neves, G., Riccio, F., Machado, C.B., Cheesbrough, A., R’Bibo, L., Burrone, J., and Lieberam, I. (2023). Aberrant axon initial segment plasticity and intrinsic excitability of ALS hiPSC motor neurons. Cell reports 42. 10.1016/j.celrep.2023.113509.

22. Paulsen, B., Velasco, S., Kedaigle, A.J., Pigoni, M., Quadrato, G., Deo, A.J., Adiconis, X., Uzquiano, A., Sartore, R., Yang, S.M., et al. (2022). Autism genes converge on asynchronous development of shared neuron classes. Nature 602, 268–273. 10.1038/s41586-021-04358-6.

23. Spratt, P.W.E., Alexander, R.P.D., Ben-Shalom, R., Sahagun, A., Kyoung, H., Keeshen, C.M., Sanders, S.J., and Bender, K.J. (2021). Paradoxical hyperexcitability from Na(V)1.2 sodium channel loss in neocortical pyramidal cells. Cell reports 36, 109483. 10.1016/j.celrep.2021.109483.

24. Quadrato, G., Nguyen, T., Macosko, E.Z., Sherwood, J.L., Min Yang, S., Berger, D.R., Maria, N., Scholvin, J., Goldman, M., Kinney, J.P., et al. (2017). Cell diversity and network dynamics in photosensitive human brain organoids. Nature 545, 48–53. 10.1038/nature22047.

25. Fair, S.R., Julian, D., Hartlaub, A.M., Pusuluri, S.T., Malik, G., Summerfied, T.L., Zhao, G., Hester, A.B., Ackerman, W.E., Hollingsworth, E.W., et al. (2020). Electrophysiological Maturation of Cerebral Organoids Correlates with Dynamic Morphological and Cellular Development. Stem Cell Reports 15, 855–868. 10.1016/j.stemcr.2020.08.017.

26. Paşca, A.M., Sloan, S.A., Clarke, L.E., Tian, Y., Makinson, C.D., Huber, N., Kim, C.H., Park, J.-Y., O’Rourke, N.A., Nguyen, K.D., et al. (2015). Functional cortical neurons and astrocytes from human pluripotent stem cells in 3D culture. Nature Methods 12, 671–678. 10.1038/nmeth.3415.

27. Tamura, S., Nelson, A.D., Spratt, P.W.E., Kyoung, H., Zhou, X., Li, Z., Zhao, J., Holden, S.S., Sahagun, A., Keeshen, C.M., et al. (2022). CRISPR activation rescues abnormalities in *SCN2A* haploinsufficiency-associated autism spectrum disorder. bioRxiv, 2022.2003.2030.486483. 10.1101/2022.03.30.486483.

28. Nunes, C., Gorczyca, G., Mendoza-deGyves, E., Ponti, J., Bogni, A., Carpi, D., Bal-Price, A., and Pistollato, F. (2022). Upscaling biological complexity to boost neuronal and oligodendroglia maturation and improve in vitro developmental neurotoxicity (DNT) evaluation. Reproductive Toxicology 110, 124–140. 10.1016/j.reprotox.2022.03.017.

29. Zhang, J., Eaton, M., Chen, X., Zhao, Y., Kant, S., Deming, B.A., Harish, K., Nguyen, H.P., Shu, Y., Lai, S., et al. (2025). Restoration of excitation/inhibition balance enhances neuronal signal-to-noise ratio and rescues social deficits in autism-associated Scn2a-deficiency. bioRxiv, 2025.2003.2004.641498. 10.1101/2025.03.04.641498.

30. Sanders, S.J., Campbell, A.J., Cottrell, J.R., Moller, R.S., Wagner, F.F., Auldridge, A.L., Bernier, R.A., Catterall, W.A., Chung, W.K., Empfield, J.R., et al. (2018). Progress in Understanding and Treating SCN2A-Mediated Disorders. Trends in neurosciences 41, 442–456. 10.1016/j.tins.2018.03.011.

31. Epifanio, R., Giorda, R., Merlano, M.C., Zanotta, N., Romaniello, R., Marelli, S., Russo, S., Cogliati, F., Bassi, M.T., and Zucca, C. (2021). SCN2A Pathogenic Variants and Epilepsy: Heterogeneous Clinical, Genetic and Diagnostic Features. Brain Sci 12. 10.3390/brainsci12010018.

32. Carvill, G.L., Matheny, T., Hesselberth, J., and Demarest, S. (2021). Haploinsufficiency, Dominant Negative, and Gain-of-Function Mechanisms in Epilepsy: Matching Therapeutic Approach to the Pathophysiology. Neurotherapeutics 18, 1500–1514. 10.1007/s13311-021-01137-z.

33. Gao, Y., Shonai, D., Trn, M., Zhao, J., Soderblom, E.J., Garcia-Moreno, S.A., Gersbach, C.A., Wetsel, W.C., Dawson, G., Velmeshev, D., et al. (2024). Proximity analysis of native proteomes reveals phenotypic modifiers in a mouse model of autism and related neurodevelopmental conditions. Nature communications 15, 6801. 10.1038/s41467-024-51037-x.

34. Skarnes, W.C., Pellegrino, E., and McDonough, J.A. (2019). Improving homology-directed repair efficiency in human stem cells. Methods 164-165, 18–28. 10.1016/j.ymeth.2019.06.016.

35. Skarnes, W.C., Ning, G., Giansiracusa, S., Cruz, A.S., Blauwendraat, C., Saavedra, B., Holden, K., Cookson, M.R., Ward, M.E., and McDonough, J.A. (2021). Controlling homology-directed repair outcomes in human stem cells with dCas9. bioRxiv, 2021.2012.2016.472942. 10.1101/2021.12.16.472942.

36. Pantazis, C.B., Yang, A., Lara, E., McDonough, J.A., Blauwendraat, C., Peng, L., Oguro, H., Kanaujiya, J., Zou, J., Sebesta, D., et al. (2022). A reference human induced pluripotent stem cell line for large-scale collaborative studies. Cell Stem Cell 29, 1685– 1702 e1622. 10.1016/j.stem.2022.11.004.

37. Pasca, S.P., Arlotta, P., Bateup, H.S., Camp, J.G., Cappello, S., Gage, F.H., Knoblich, J.A., Kriegstein, A.R., Lancaster, M.A., Ming, G.L., et al. (2025). A framework for neural organoids, assembloids and transplantation studies. Nature 639, 315–320. 10.1038/s41586-024-08487-6.

38. Zhang, J., Zhang, C., Chen, X., Wang, B., Ma, W., Yang, Y., Zheng, R., and Huang, Z. (2021). PKA-RIIβ autophosphorylation modulates PKA activity and seizure phenotypes in mice. Communications Biology 4, 263. 10.1038/s42003-021-01748-4.

39. Wang, J., He, X., Meng, H., Li, Y., Dmitriev, P., Tian, F., Page, J.C., Lu, Q.R., and He, Z. (2020). Robust Myelination of Regenerated Axons Induced by Combined Manipulations of GPR17 and Microglia. Neuron 108, 876–886.e874.

40. Chen, S., Zhou, Y., Chen, Y., and Gu, J. (2018). fastp: an ultra-fast all-in-one FASTQ preprocessor. Bioinformatics 34, i884–i890. 10.1093/bioinformatics/bty560.

41. Dobin, A., Davis, C.A., Schlesinger, F., Drenkow, J., Zaleski, C., Jha, S., Batut, P., Chaisson, M., and Gingeras, T.R. (2013). STAR: ultrafast universal RNA-seq aligner. Bioinformatics 29, 15–21. 10.1093/bioinformatics/bts635.

42. McKenna, A., Hanna, M., Banks, E., Sivachenko, A., Cibulskis, K., Kernytsky, A., Garimella, K., Altshuler, D., Gabriel, S., Daly, M., and DePristo, M.A. (2010). The Genome Analysis Toolkit: a MapReduce framework for analyzing next-generation DNA sequencing data. Genome Res 20, 1297–1303. 10.1101/gr.107524.110.

43. Brouard, J.S., Schenkel, F., Marete, A., and Bissonnette, N. (2019). The GATK joint genotyping workflow is appropriate for calling variants in RNA-seq experiments. J Anim Sci Biotechnol 10, 44. 10.1186/s40104-019-0359-0.

44. Liao, Y., Smyth, G.K., and Shi, W. (2014). featureCounts: an efficient general purpose program for assigning sequence reads to genomic features. Bioinformatics 30, 923–930. 10.1093/bioinformatics/btt656.

45. Love, M.I., Huber, W., and Anders, S. (2014). Moderated estimation of fold change and dispersion for RNA-seq data with DESeq2. Genome Biol 15, 550. 10.1186/s13059-014-0550-8.

46. Ihaka, R., and Gentleman, R. (1996). R: a language for data analysis and graphics. Journal of computational and graphical statistics 5, 299–314.

47. Risso, D., Ngai, J., Speed, T.P., and Dudoit, S. (2014). Normalization of RNA-seq data using factor analysis of control genes or samples. Nat Biotechnol 32, 896–902. 10.1038/nbt.2931.

48. Robinson, M.D., McCarthy, D.J., and Smyth, G.K. (2010). edgeR: a Bioconductor package for differential expression analysis of digital gene expression data. Bioinformatics 26, 139–140. 10.1093/bioinformatics/btp616.

49. Yu, G., Wang, L.G., Han, Y., and He, Q.Y. (2012). clusterProfiler: an R package for comparing biological themes among gene clusters. Omics 16, 284–287. 10.1089/omi.2011.0118.

50. Krämer, A., Green, J., Pollard, J., Jr., and Tugendreich, S. (2014). Causal analysis approaches in Ingenuity Pathway Analysis. Bioinformatics 30, 523–530. 10.1093/bioinformatics/btt703.

51. Gu, Z. (2022). Complex heatmap visualization. Imeta 1, e43. 10.1002/imt2.43.

52. Blighe, K., Rana, S., and Lewis, M. (2019). EnhancedVolcano: Publication-ready volcano plots with enhanced colouring and labeling. R package version 1.2. 0. R package version 1.16.

